# High-throughput genetic mapping discovers novel zinc toxicity response loci in *Drosophila melanogaster*

**DOI:** 10.1101/2025.06.12.659401

**Authors:** Katherine M. Hanson, Anthony D. Long, Stuart J. Macdonald

## Abstract

Heavy metals are a widespread environmental contaminant, and even low levels of some metals can disrupt cellular processes and result in DNA damage. However, the consequences of metal exposure are variable among individuals, with susceptibility to metal toxicity representing a complex trait influenced by genetic and non-genetic factors. To uncover toxicity response genes, and better understand responses to metal toxicity, we sought to dissect resistance to zinc, a metal required for normal cellular function, which can be toxic at high doses. To facilitate efficient, powerful discovery of Quantitative Trait Loci (QTL) we employed extreme, or X-QTL mapping, leveraging a multiparental, recombinant *Drosophila melanogaster* population. Our approach involved bulk selection of zinc-resistant individuals, sequencing several replicate pools of selected and control animals, and identified QTL as genomic positions showing consistent allele frequency shifts between treatments. We successfully identified seven regions segregating for resistance/susceptibility alleles, and implicated several strong candidate genes. Phenotypic characterization of populations derived from selected or control animals revealed that our selection procedure resulted in greater egg-to-adult emergence, and a reduced developmental delay on zinc media. We subsequently measured emergence and development time for a series of midgut-specific RNAi gene knockdowns and matched genetic controls raised in both zinc-supplemented and normal media. This identified ten genes with significant genotype-by-treatment effects, including *pHCl-2*, which encodes a zinc sensor protein. Our work highlights recognized and novel contributors to zinc toxicity resistance in flies, and provides a pathway to a broader understanding of the biological impact of metal toxicity.

**ARTICLE SUMMARY:** Starting with an outbred *Drosophila melanogaster* population we repeatedly selected for groups of individuals showing high resistance to toxic levels of zinc during development. Pooled sequencing of these groups, along with matched groups of control individuals, enabled the identification of seven genomic regions – or QTL – contributing to zinc toxicity resistance. Midgut-specific RNAi of genes implicated by these QTL yielded ten genes impacting developmental traits in zinc-supplemented media, including *MTF-1* (a metal response transcription factor) and *pHCl-2* (a zinc sensor protein).

## INTRODUCTION

Human exposure to toxic levels of various heavy metals has risen due to environmental contamination resulting from mining operations, and the broad use of metal compounds in a range of agricultural and industrial sectors (Jaishankar et al., 2014; Karri et al., 2016; Kim et al., 2019; Rehman et al., 2018; Tchounwou et al., 2012). Exposure to heavy metals primarily occurs through ingestion of contaminated water or food grown in contaminated soil, but can additionally occur via inhalation (Rehman et al., 2018; Tchounwou et al., 2012). Exposure to a high level of heavy metals can harm cells by inducing the production of reactive oxygen species which can damage DNA through oxidative stress, alter the balance of essential elements (for instance, where lead ions can replace iron ions), and affect enzymatic activity (Fu & Xi, 2020; Jaishankar et al., 2014; Rehman et al., 2018). Such metal-induced change can yield severe health consequences, increasing the risk for cancer, cognitive dysfunction, and organ damage (Goutam Mukherjee et al., 2022; Karri et al., 2016; Rehman et al., 2018; Tchounwou et al., 2012). Given the pervasive negative consequences of heavy metal exposure, it is critical to understand how organisms physiologically respond to metal ingestion, and the molecular pathways that are involved.

Although many heavy metals – for instance, cadmium and lead – have no physiological role, metals such as copper, iron, manganese, and zinc are essential heavy metals required for a plethora of biological functions (Kim et al., 2019; Tchounwou et al., 2012). Zinc is abundant in living organisms, has a variety of roles as both a structural and catalytic component of proteins (Kambe et al., 2015), is involved in cell signaling (Fukada et al., 2011), and is estimated to be able to bind to 10% of the human proteome (Andreini et al., 2006). Given its key role in many processes, maintaining an appropriate cellular level of zinc is critical for many biological functions: Zinc deficiency can lead to a range of health problems (Fischer Walker et al., 2009; Prasad, 2013), and conversely excess consumption of zinc – such as the overuse of zinc dietary supplements (Irving et al., 2003; Willis et al., 2005), denture cream (Lasater, 2017; Willis et al., 2005), or inhaling zinc-containing fumes from industrial process such as welding (ATSDR, 2005; Cooper, 2008; Noel & Ruthman, 1988) – can lead to toxicity.

Toxic levels of zinc can induce oxidative stress, inhibit mitochondrial function leading to reduced ATP levels, and induce cell death (Lemire et al., 2008; Maret, 2008; Sensi et al., 2009). Additionally, an important consequence of zinc toxicity is copper deficiency whose symptoms include anemia, leukopenia and neutropenia (Agnew & Slesinger, 2020; Hoffman et al., 1988; Willis et al., 2005). This reduction in copper level is likely due to the high levels of zinc inducing the expression of metallothionein (MT) proteins (a protein family important in metal detoxification), which have a greater affinity for copper than for zinc, resulting in lower levels of free copper (Agnew & Slesinger, 2020; Harold, 2014; Navarro & Schneuwly, 2017; Yiwen et al., 2022).

Central to zinc homeostasis is *MTF-1* (*metal transcription factor 1*), a zinc-sensing transcription factor. When cellular zinc levels are high, MTF-1 will increase the expression of genes regulated by the metal response element enhancer (Andrews, 2001; Stuart et al., 1985; B. Zhang et al., 2001). Such genes include MTs (Egli et al., 2003; B. Zhang et al., 2001), along with some transporters belonging to the zinc transporter (ZnT) family (Guo et al., 2010; Langmade et al., 2000; Yepiskoposyan et al., 2006) which are responsible for transporting cytosolic zinc out to the extracellular matrix or into intracellular compartments. Additionally, MTF-1 will repress expression of some proteins belonging to the zinc import protein (ZIP) family (Lichten et al., 2011; Zheng et al., 2008) which are responsible for importing zinc into the cytosol.

Many of the proteins that regulate zinc are evolutionarily conserved between mammals and *Drosophila melanogaster* (Lye et al., 2012; B. Zhang et al., 2001), making flies an excellent model to study zinc homeostasis and toxicity. For example, adding the human MTF-1 sequence into *MTF-1* null *Drosophila* cell lines rescues sensitivity to heavy metals, while the *Drosophila* version of MTF-1 is able to rescue sensitivity in MTF-1 knockout mouse cells (Balamurugan et al., 2004). Additionally, the *Drosophila* genome encodes 7 ZnT proteins and 10 ZIP proteins that are homologous to mammalian transporters (Calap-Quintana et al., 2017; Xiao & Zhou, 2016), and flies also possess 6 MTs which help protect against metal toxicity (Luo et al., 2020).

A great deal has been learned about the biological roles of zinc, and the genes, pathways, and mechanisms involved in maintaining zinc homeostasis and avoiding toxicity, from functional studies in whole animals that rely on gene knockouts (Egli et al., 2003; Palmiter & Findley, 1995; Yepiskoposyan et al., 2006), RNAi-based gene expression knockdowns (Lye et al., 2012; B. Zhang et al., 2001), or overexpression of critical genes (Gaither & Eide, 2001; Lye et al., 2012), as well as from cell-based functional genomic screens (Mohr et al., 2018). However, such studies typically employ an esoteric (and often limited) set of genetic backgrounds, which might make inferring the impact of mutations difficult to generalize (Chandler et al., 2013; Sittig et al., 2016). Since there is population-level variation for responses to the toxic effects of heavy metals in general (Evans et al., 2018; Warrington et al., 2015; Xenakis et al., 2022; Zhou et al., 2017), and zinc specifically (Evans et al., 2020), isolating and characterizing loci that segregate for allelic variation impacting susceptibility is a complementary route to understanding zinc biology. For instance, studies in rice (J. Zhang et al., 2017), *Arabidopsis helleri* (Willems et al., 2007), and *Caenorhabditis elegans* (Evans et al., 2020) have isolated QTL (Quantitative Trait Loci) that contribute to variation in zinc resistance, and have led to the identification of novel loci underlying metal resistance/susceptibility.

Here we seek to further understand the biology of zinc homeostasis and toxicity by genetically dissecting variation in zinc resistance using *D. melanogaster*. We take advantage of the “extreme QTL” or X-QTL mapping framework that is an elaboration of bulked segregant analysis (Michelmore et al., 1991). Initially employed in yeast (Ehrenreich et al., 2010), X-QTL mapping starts with a large recombinant population and bulk selects for a phenotype of interest. Individuals who pass the selection threshold – in the present case, adults that emerge from embryos raised through media containing a high level of zinc – are pooled, sequenced, and compared to a pool of non-selected, control animals. QTL are identified as genomic positions showing consistent differences in allele frequency between several replicate pairs of selected and control pools. The X-QTL design we employ extends the approach by starting from a highly-recombinant, outbred multiparental base population derived from the *Drosophila* Synthetic Population Resource (DSPR) (King, Merkes, et al., 2012) and enables powerful QTL mapping, and excellent mapping resolution (Macdonald et al., 2022). The X-QTL approach is particularly amenable to dissecting toxicity traits since bulk phenotypic selection for such traits is straightforward and can be very efficient. For instance, we have successfully used X-QTL mapping to identify loci underlying resistance to caffeine (Macdonald et al., 2022) and the insecticide malathion (Macdonald & Long, 2022). Although these previous experiments discovered novel QTL associated with trait variation, simulations suggest power was limited by a lack of replication and the modest numbers of individuals assayed (Macdonald et al., 2022). Therefore, in the present study we employed 12 experimental replicates, testing tens of thousands of individuals, allowing us to isolate 7 QTL that impact developmental zinc resistance. Follow-up midgut RNAi of a subset of genes within QTL intervals implicated several candidates in the control of trait variation, some of which represent novel contributors to zinc toxicity tolerance in flies, including *pHCl-2* (pH-sensitive chloride channel 2) and *Ndae1* (Na^+^-driven anion exchanger 1).

## MATERIALS & METHODS

### X-QTL base population

We employed the same DSPR-derived base population used in two previous X-QTL studies (Macdonald et al., 2022; Macdonald & Long, 2022). Briefly, the DSPR is a set of multiparental advanced generation intercross RILs (Recombinant Inbred Lines) derived from 8 inbred founder strains (King, Merkes, et al., 2012). We collected 10 eggs from each of 663 DSPR population A RILs, raised them in standard *Drosophila* media bottles, and placed adult progeny in a population cage. The population was provisioned weekly with fresh media in bottles, and maintained for 64 weeks before egg collection for the first replicate of the zinc selection X-QTL assay (see below). Assuming the population experienced one generation every two weeks, we estimate that the 12 X-QTL replicates were executed 31-36 generations following base population founding.

### Zinc selection X-QTL assay

Each of the 12 replicates of the zinc selection assay was executed as follows. Several apple juice agar plates, each holding a small amount of live yeast paste in the center, were placed in the base population cage for ∼24-hours. Subsequently, plates were filled with 1X phosphate-buffered saline (PBS), eggs were gently displaced from the plate surface with a paintbrush, and the egg suspension poured into a 50-ml conical tube. Eggs were rinsed with 1X PBS to remove any media residue, and the majority of the 1X PBS drained from the clean eggs, leaving them just submerged. We then pipetted eggs into a series of 6-oz *Drosophila* bottles (Fisher Scientific, AS-355) using a wide-bore pipette tip. The media in each bottle contained 9-g Instant *Drosophila* Medium (Carolina Biological Supply Company, Formula 4-24) reconstituted with 40-ml of either water or a 25-mM zinc chloride solution (ZnCl_2_, CAS Number 7646-85-7, Millipore-Sigma, 793523). We pipetted 24-µl eggs into each water bottle (the non-selection, control treatment), and 48-µl eggs into each zinc bottle (the selection treatment). The greater number of embryos per zinc bottle limited the total number of bottles required, while the larval/adult density per bottle remained low given the toxicity of the zinc chloride concentration employed. For each replicate, we set up four water control bottles and between 8 and 21 zinc bottles (see Supplementary Table 1).

Bottles were checked for emerged adults every 1-2 days. Flies emerged from the water bottles on days 10-15 following setup, while they emerged from the zinc bottles on days 15-22 due to the developmental delay caused by the zinc treatment. Emerged adults were counted, separated by sex and treatment, and females were frozen at −20°C for subsequent DNA isolation. For replicates R9 and R10, flies were not immediately frozen, and instead were allowed to lay eggs to derive subpopulations to enable a more detailed examination of the phenotypic effects of selection (see below).

### Pooled DNA isolation

We routinely collected more non-selected, control females from the water bottles than selected females from the zinc bottles. To maintain the same pool size for the two treatments within each replicate we randomly chose a subset of the non-selected, control females to match the actual number of zinc-selected females that emerged. For five replicates (R5, R6, R8, R11, R12) we had enough control females to generate two independent control pools, enabling a direct examination of variation among pools of control animals. Each of the resulting 29 pools of flies were homogenized in bulk, and DNA was isolated from each using the Puregene Cell Kit (Qiagen, 158046) with a modified protocol (Supplementary Text 1). Resuspended DNA was quantified using a Qubit fluorometer (ThermoFisher).

### DNA library preparation and sequencing

Each of the 29 pooled DNA samples was diluted to 450-ng in 30-µl, and libraries were constructed following the Illumina DNA Prep protocol (Illumina, 20018705) with unique dual indices (Illumina, 20027213). The 29 libraries ranged in fragment size from 403-bp to 577-bp (Agilent TapeStation 4150), were pooled at equal concentrations, and sequenced with PE150 reads on an Illumina NovaSeq 6000 SP flowcell at the University of Kansas Medical Center Genome Sequencing Facility.

### Haplotype estimation and QTL identification

QTL were identified following a similar framework as used in a previous X-QTL study employing the same base population (Macdonald & Long, 2022). Briefly, raw reads from each of the 29 X-QTL samples, along with reads from the founder strains of DSPR population A (King, Merkes, et al., 2012), were mapped to Release 6 (dm6) of the *D. melanogaster* reference genome with bwa-mem (Li, 2013). We used bcftools (Li, 2011) to obtain REF and ALT counts for every SNP, and these were converted to REF allele frequencies at each SNP for each sample. We subsequently moved along the genome in overlapping 1.5-cM windows, and for each window for each X-QTL sample used the R/limSolve package (Soetaert et al., 2009) to estimate the founder haplotype frequencies based on the observed SNP frequencies in the X-QTL sample, and the known founder haplotypes. For additional detail of the method, along with validation of the accuracy of this estimation approach, see both Linder et al. 2020 and Macdonald et al. 2022.

The estimated founder haplotype frequencies for each replicate and treatment were arcsine square-root transformed (ASF), then tested for differentiation between treatments using an ANOVA (*ASF* ∼ *Treatment* + *Haplotype* + *Treatment* × *Haplotype*). For each window we calculated a −log_10_(*P*) value by testing the *Treatment* × *Haplotype* interaction term against *Replicate* × *Treatment* × *Haplotype* as the error term. Subsequently we LOESS smoothed the genomewide set of −log_10_(*P*) values to compensate for window-to-window variation, identified QTL peaks as those positions exceeding a significance threshold of −log_10_(*P*) = 4, and for each peak identified the confidence interval on location as a 3 −log_10_(*P*) drop confidence interval.

### Phenotypic consequences of selection for increased developmental zinc resistance

We generated several populations/strains, and executed several experiments to examine the phenotypic impact of zinc exposure during development.

#### Deriving subpopulations from emerged X-QTL animals

For replicates R9 and R10 of the X-QTL study, we collected all emerging flies from the two treatment regimes as living animals. All those adults emerging from the zinc-selection treatment for a given replicate were split among several vials containing cornmeal-yeast-molasses media, and allowed to lay eggs for 72 hours. Flies emerging from the non-selected, water control treatment were treated similarly, except we only used a random subset of the animals to match the (lower) number of animals emerging from the zinc selection treatment. The four subpopulations – R9C, R9Z, R10C, R10Z (where “C” and “Z” represent the control and zinc treatments, respectively) – were maintained in sets of vials for several generations and used for phenotyping (below).

#### Generating lines from X-QTL subpopulations

In an effort to stabilize the genotypes of the outbred subpopulations, following seven generations of subpopulation maintenance, we established 22 lines from R9C (R9C-lines) and 23 from R9Z (R9Z-lines) using single, mated females from the subpopulations, and followed this with 6-8 generations of inbreeding (crosses between a single virgin female and one male). Clearly these lines are only partially inbred, and to reflect this we refer to them as “semi-inbred lines” throughout.

#### Establishing new DSPR-derived cohorts of zinc-selected and control animals

Part way through the project we disposed of the X-QTL base population. To continue our phenotypic testing, we derived a new population by collecting 10 eggs from each of 83 DSPR population A RILs, allowing the population to mix/recombine for five generations. Next, we collected eggs and followed the same protocol described above (see the “Zinc selection X-QTL assay” section) yielding cohorts of zinc-selected and non-selected, control animals. These groups were allowed to mate and lay eggs, were maintained for two generations, and then were tested in phenotypic assays (see below). We refer to these as the selected and non-selected phenotyping-only test cohorts.

#### Assaying developmental resistance to heavy metals

Test vials contained 50 eggs on media containing 1.8-g Instant *Drosophila* Medium mixed with either 8-ml of water or a heavy metal salt solution. We used solutions of zinc chloride (10- or 25-mM depending on the experiment), 2-mM copper sulfate (CuSO_4_, CAS number 7758-98-7, Millipore-Sigma, C1297), and 0.2-mM cadmium chloride (CdCl_2_, CAS number 10108-64-2, Millipore-Sigma, 655198). Emerged flies were sexed/counted up to 30 days following egg collection. We calculated the fraction of adults emerging per vial, and the egg-to-adult development time (in days) for every individual emerged adult.

We assayed developmental zinc resistance in all four subpopulations derived from replicates R9 and R10 of the X-QTL study (R9C, R9Z, R10C, R10Z) by testing 10 25-mM ZnCl_2_ vials and 2-3 water vials per subpopulation. The two sets of semi-inbred lines (the R9C- and R9Z-lines) were tested for developmental resistance to a lower concentration of zinc (∼3 10-mM ZnCl_2_ vials per line, and 63-68 per set), copper (1 vial/line, 22-23 vials/set), and cadmium (1 vial/line, 22-23 vials/set), with water controls (1 vial/line, 22-23 vials/set). We were forced to reduce the test ZnCl_2_ concentration to 10-mM given that pilot work showed most semi-inbred lines had zero emergence at 25-mM ZnCl_2_. Finally, we tested eggs from the selected and non-selected phenotyping-only test cohorts for resistance to zinc (40 25-mM ZnCl_2_ vials per cohort), copper (40 vials per cohort), and cadmium (40 vials per cohort), along with 38-39 water vials per cohort.

#### Measuring zinc resistance in adults

Assays were carried out in standard narrow fly vials containing 1.8-g Instant *Drosophila* Medium with 8-ml of either water or 100-mM ZnCl_2_. Twenty 2-6 day old, likely mated females were placed into each vial, and dead flies were counted daily until day 7 (by this point all flies in the zinc-containing vials had died). Flies in water-control vials were moved to fresh vials every 2-3 days to maintain media quality (given the absence of a toxic level of zinc, the action of larvae made the instant media wet/sticky). We assayed adult zinc resistance in the four X-QTL subpopulations (R9Z, R9C, R10Z, R10C) over two batches, using 10 zinc and 2 water vials per subpopulation in batch 1, and 20 zinc and 2 water vials per subpopulation in batch 2. We also assayed 25 zinc vials and 6 water vials for each of the selected and non-selected phenotyping-only test cohorts.

### Testing X-QTL candidate genes via RNAi

Collectively, the X-QTL intervals we resolved contain several hundred protein-coding genes. Candidate genes for RNAi testing were prioritized using the following information. First, we identified those genes known to be involved in zinc homeostasis (i.e., *MTF-1*, and any members of the MT, ZnT or ZIP families). Second, we examined FlyBase (release FB2022_03) information associated with each gene (Gramates et al., 2022), particularly focusing on genes associated with the gene ontology (GO) terms zinc ion binding (GO:0008270), intracellular zinc homeostasis (GO:0006882), response to oxidative stress (GO:0006979), as well as other terms related to zinc and other essential metals (i.e., magnesium, copper, cadmium, and iron). Third, we cross-referenced genes under our mapped QTL with genes identified in a cell-based RNAi screen for zinc toxicity modifiers, and those that showed a significant transcriptional change in response to zinc exposure (Mohr et al., 2018, see their File S1 and S2 datasets). Fourth, we marked genes found in previous metal toxicity studies and relevant review articles (Navarro & Schneuwly, 2017; Redhai et al., 2020). Collectively, we identified 24 candidate genes within our mapped QTL (Supplementary Table 2). We note that ongoing optimization of our X-QTL mapping routines throughout this project led to slight differences in the locations of QTL boundaries between the time we evaluated candidate genes, and the time of writing, so 3/24 candidates are just outside of the X-QTL boundaries we report here (Table 1, Supplementary Table 2).

**Table 1.** Mapped zinc resistance QTL.

| QTL | Chromosome | Peak Position (bp) <sup>a</sup> | Physical Interval (bp) <sup>a</sup> | QTL Size (Kb) | QTL Size (cM) | Genes <sup>b</sup> | Candidate Genes |
| --- | --- | --- | --- | --- | --- | --- | --- |
| A | X | 13,945,053 | 13,796,451 – 14,093,253 | 300 | 1.30 | 21 | *ben* |
| B | 2L | 7,135,043 | 6,829,342 – 7,325,061 | 560 | 2.20 | 58 | *Gart*<sup>c</sup>, *Mnn1*, *CG4496*, *Ndae1* |
| C | 2R | 15,413,541 | 15,116,465 – 15,886,141 | 780 | 2.85 | 88 | *Trpm*, *Vha36-1*<sup>c</sup>, *Vha14-1*<sup>c</sup> |
| D | 3L | 8,925,893 | 8,352,067 – 9,512,583 | 1160 | 4.70 | 166 | *GluRIB*, *Hsp22*, *MTF-1* |
| E | 3R | 18,864,745 | 18,717,419 – 18,987,824 | 280 | 0.65 | 30 | *CG14302*<sup>c</sup>, *Mekk1*, *Octa2R*, *Xrp1*, *ATPsynD*<sup>d</sup> |
| F | 3R | 23,906,956 | 23,458,263 – 24,716,284 | 1250 | 4.50 | 190 | *RanBP3*, *Nup98-96*, *Ndc1* |
| G | 3R | 30,848,984 | 30,368,845 – 31,020,051 | 820 | 0.80 | 78 | *Fer1HCH*, *Fer2LCH*, *dj-1β*, *pHCl-2*<sup>d</sup>, *CG11318*<sup>d</sup> |

<sup>a</sup> Physical positions are determined from the corresponding genetic positions, and are based on Release 6 of the *D. melanogaster* reference genome. Intervals are defined as 3 $-\log_{10}(P)$ drops from the QTL peak LOD score position.
<sup>b</sup> The number of annotated protein-coding genes. Annotations sourced from Flybase release FB2022\_06 (Gramates et al., 2022).
<sup>c</sup> These genes did not have TRiP UAS-RNAi strains available from the BDSC so were not tested via RNAi.
<sup>d</sup> Genes not within the final set of X-QTL intervals. When genes were selected for functional testing a slightly different analysis scheme was being employed to that reported here. *ATPsynD* is within 120-kb of the left end of the QTL-E interval, and *CG11318* and *pHCl-2* are within 90- kb and 290-kb of the left end of the QTL-G interval, respectively. We note that QTL-G is at the terminal end of chromosome 3R (Figure 1).

We knocked down candidate genes using the X-linked *mex1*-GAL4 driver (Bloomington *Drosophila* Stock Center, BDSC 91367), which is reported to express GAL4 in the middle midgut from embryos through to second instar larvae (Phillips & Thomas, 2006). All UAS-RNAi and control stocks were derived from the Transgenic RNAi Project (TRiP) resource (Perkins et al., 2015). We used the following 4 control strains: UAS-GFP (BDSC 35786), UAS-Luciferase (35788), empty attP2 docking site (36303), and an empty attP40 docking site (36304). The following UAS-RNAi lines were used to successfully test 18 candidate genes: *ATPsynD* (33740), *bendless* (28721), *CG11318* (51792), *CG4496* (51428), *dj-1β* (38999), *GluRIB* (67843), *Hsp22* (41709, which will also impact expression of the overlapping gene *CG4456*), *Mekk1* (28587), *Mnn1* (51862), *MTF-1* (33381 and 34094), *Ndae1* (62177), *Ndc1* (67275), *Nup98-96* (28562), *Octα2R* (50678), *pHCl-2* (26003), *RanBP3* (40948), *Trpm* (51713 and 57871), and *Xrp1* (34521 and 51054). Initial tests of RNAi for two candidate genes – *Fer1HCH* (60000) and *Fer2LCH* (44067) – revealed almost no adult GAL4:UAS-RNAi flies emerging on control media, suggesting knockdown of these genes via *mex1*-GAL4 leads to lethality in the absence of zinc supplementation; we did not pursue these genes further. Finally, we did not test four of our candidate genes – *Gart*, *Vha36-1*, *Vha14-1*, *CG14302* – since TRiP RNAi lines were not available.

Virgin female *mex1*-GAL4 females were crossed to males from the RNAi control and UAS-RNAi strains, avoiding balancer-containing males for those strains segregating for a balancer chromosome. After the first day of mating in standard media vials, crosses were tipped into custom egg laying chambers. Each chamber is constructed from a polypropylene fly vial (Fisher Scientific, AS507) with the base removed and replaced with a cotton ball. Flies are enclosed in the chamber by capping it with a polyethylene flanged plastic cap (MOCAP, FCS13/16NA1) containing ∼4.5-ml of cornmeal-yeast-molasses media plus a small amount of live yeast paste to encourage egg laying. Media caps were replaced daily for 4 days, and each day eggs were collected manually. Eggs were placed in groups of 50 into water-control or 10-mM ZnCl_2_ treatment vials (standard narrow fly vials containing 1.8-g Instant *Drosophila* Medium reconstituted with 8-ml of solution). Following egg collection, we sexed/counted emerged flies for 30 days. Five batches of the experiment were completed, resulting in 17-21 replicate vials per treatment per candidate gene RNAi, and 28-29 replicate vials per treatment for each RNAi control genotype. As for the phenotyping populations and strains (above) we measured adult emergence on a per-vial basis, while egg-to-adult development time was assessed for every emerging adult in days.

We note that the RNAi experiment used a lower ZnCl_2_ concentration than our X-QTL experiment (10-mM *versus* 25-mM); This was enforced by pilot RNAi experiments using the higher concentration, during which no eggs developed into adults.

### Fly maintenance

Aside from the X-QTL base population and the second population created from 83 RILs, (which were both maintained in population cages at normal laboratory conditions), all flies were reared/tested in an incubator at 25°C, 50% humidity, with a 12-hour dark:light cycle. For normal development/maintenance, all populations and strains were maintained using a cornmeal-yeast-molasses media (see Supplementary Text 2).

### Data availability

The DSPR RILs used in the construction of the X-QTL base population are available from the BDSC. Raw X-QTL sample FASTQ sequencing data is available from the NCBI SRA under accession number PRJNA1127662. All summary information and statistical results are presented in supplementary files accompanying this work, and all analysis code is available on GitHub (https://github.com/Hanson19/Zinc-X-QTL).

## RESULTS

### Discovery of zinc toxicity resistance loci

We established an outbred base population by mixing together animals from >600 8-way population A DSPR RILs (King, Macdonald, et al., 2012) resulting in a population that segregates for the 8 DSPR founder haplotypes at any given genomic position. We then created a developmental zinc toxicity assay that can be easily carried out in bulk at large scale, and selected animals with genotypes that confer relatively greater resistance to zinc. Over 12 replicates of this X-QTL design, eggs were collected from the base population, and allowed to develop to adulthood on either a non-selection, control treatment media or a selection treatment media containing 25mM ZnCl_2_. We did not count the number of embryos tested, but based on the number of flies emerging from the non-selection, control treatment, we estimate that each replicate tested around 4,300 female embryos on the zinc treatment (range = 2,600-6,000), and that ∼7% of these survived to adulthood (Supplementary Table 1). For each replicate, we isolated DNA from a pool of all females emerging from the zinc treatment, and a pool of an equivalent number of females coming from the non-selection, control treatment, and sequenced each pool to an average of ∼67X coverage.

Following read alignment and SNP calling, we moved through the genome in windows, and for each X-QTL sample estimated the frequency of the 8 DSPR founder haplotypes in that window. QTL are then detected as consistent, significant shifts in the founder haplotype frequencies between the non-selected, control samples and the zinc selection samples (see “Materials & Methods” for additional detail). We employed −log_10_(*P*)=4 as our significance threshold since our previous work showed this led to a genomewide false positive detection rate at or below 5% (Macdonald et al., 2022). Additionally, by directly comparing the five pairs of duplicate non-selection, control treatment pools (taken from replicates R5, R6, R8, R11, and R12) we see limited evidence for allele frequency shifts between duplicates, and no peaks reaching −log_10_(*P*)=4 (Supplementary Figure 1). This suggests our haplotype frequency estimates are accurate, and peaks above −log_10_(*P*)=4 likely represent true causative factors impacting zinc resistance.

We identified 7 peaks – QTL-A to QTL-G – above our significance threshold of −log_10_(*P*)=4 that are in regions of normal recombination, and these are spread across the genome on all major chromosome arms (Figure 1, Table 1). None of the QTL overlap with intervals implicated by our previous work on adult caffeine and malathion insecticide resistance using the same base population (Macdonald et al., 2022; Macdonald & Long, 2022). QTL-E does colocalize with a QTL previously identified for development viability in the presence of 2-mM copper identified using a subset of the DSPR population B RILs (Everman et al., 2021). However, we cannot know if this pair of QTL are driven by the same molecular factor since the mapping results derive from genetically distinct populations; the base population we use here is derived from the DSPR A RILs, and the A and B RILs were founded with different sets of strains (King, Merkes, et al., 2012).

**Figure 1:**
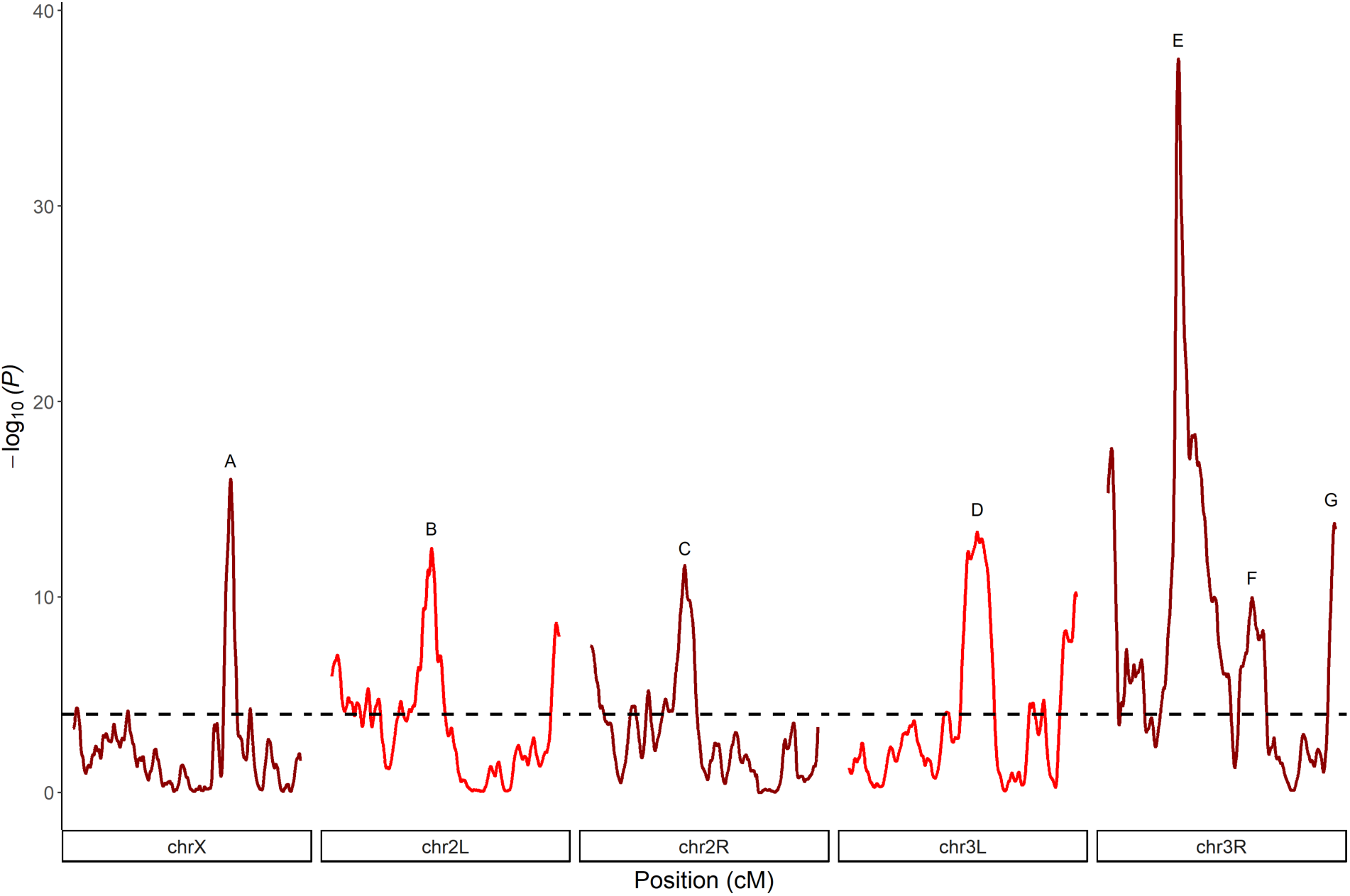
Locations of 7 mapped zinc-resistance loci. Using X-QTL mapping we identified 7 QTL – A through G – for zinc resistance throughout the genome. QTL have peaks greater than a genomewide significance threshold of −log_10_(*P*)=4 (horizontal dashed line). Additional peaks above this threshold are also apparent, but these are primarily located in centromeric regions, and while they may represent true susceptibility loci they implicate very large physical regions, and further dissection is challenging.

These 7 QTL implicate intervals of 300-1250 Kb, with the majority being less than 1000-Kb (Table 1). These intervals are smaller than in our previous DSPR-based X-QTL studies (Macdonald et al., 2022; Macdonald & Long, 2022) since we execute a greater number of experimental replicates (12 *versus* 4), which yields greater mapping power and resolution (Macdonald et al., 2022). Collectively, 631 protein-coding genes reside within our 7 QTL, and 5 QTL harbor fewer than 90 such genes. We note that our X-QTL genome scan also revealed evidence for peaks in the centromeric regions of chromosomes 2 and 3 (Figure 1). These peaks are physically large (2.3-5.5 Mb) since they implicate areas of reduced recombination, making identifying causative candidate genes extremely difficult. Thus, we focused on the 7 QTL peaks identified in regions of normal recombination.

### Haplotype frequency changes due to zinc selection at mapped QTL

Since QTL are identified as significant shifts in founder haplotype frequency due to the zinc exposure treatment, we can identify founders that possess resistant or susceptible alleles as those showing an increase or a decrease in frequency relative to controls. Figure 2 illustrates the change in founder haplotype frequency at each of the 7 peaks, showing it is routinely the case that the frequencies of only 1-2 haplotypes change substantially between the zinc and control treatments. A possible interpretation is that the functional alleles mapped via X-QTL are relatively rare, with “extreme” alleles present in just one or two founders. However, at each of the QTL we identify here, the founder haplotypes showing substantial change in frequency due to treatment are always at high frequency in the non-selection, control pools (Supplementary Figure 2). Indeed, for every one of our QTL, the frequency of the most common 1-2 founders in the control pools sums to nearly 50%.

**Figure 2:**
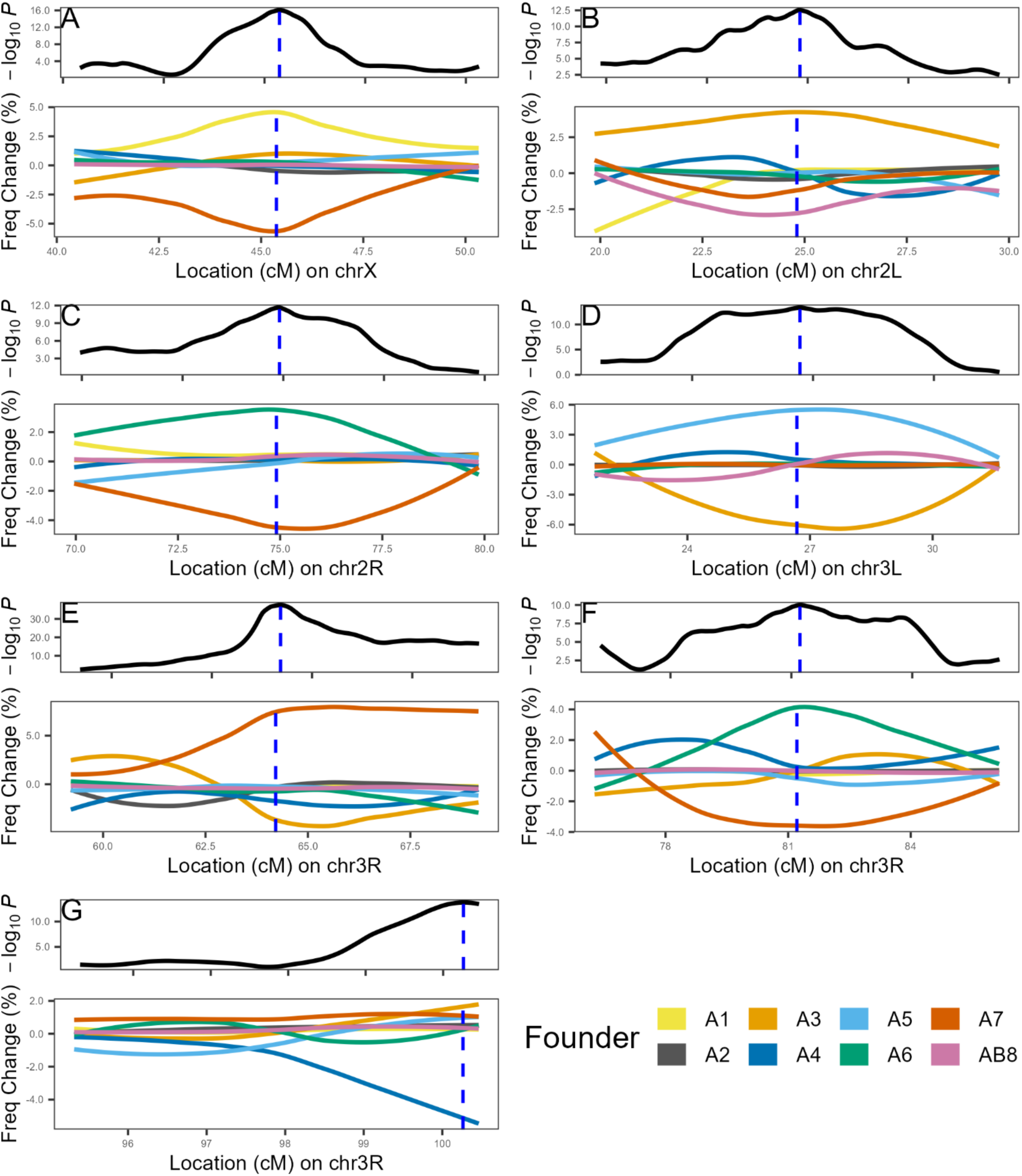
Change in founder haplotype frequencies at mapped QTL. For each of the 7 QTL (A-G) we show – top panels – a zoomed-in view of each QTL region from Figure 1, with a blue vertical dashed line highlighting the peak of the QTL. The bottom panels plot the change in the frequency of each founder haplotype between treatments (zinc-selected minus control); A frequency change above zero implies the founder is more frequent in the zinc-selected population. QTL-G is at the end of chromosome 3R, which is why the peak is on the right-hand side of the plot. (See also Supplementary Figure 2).

Unlike some other multiparental mapping panels, the DSPR RILs that we mixed to create the outbred base population exhibit considerable heterogeneity in founder haplotype frequencies along the genome, with a great deal of variation away from the 1/8 (12.5%) frequency expected (compare Figures 3-4 from Collaborative Cross Consortium, 2012 to Figure 2 from King, Merkes et al., 2012). This pattern is likely due the actions of random genetic drift and selection during the 50 generations of interbreeding that were used to build up recombination events in the populations from which the DSPR RILs were derived. As a result, since some founder haplotypes are at very low frequency at any given position in the DSPR-derived base population, our X-QTL mapping is effectively contrasting fewer than 8 alleles at any given locus. Thus, it is difficult to estimate the frequency of resistant or susceptible alleles in the species as a whole from our data.

**Figure 3:**
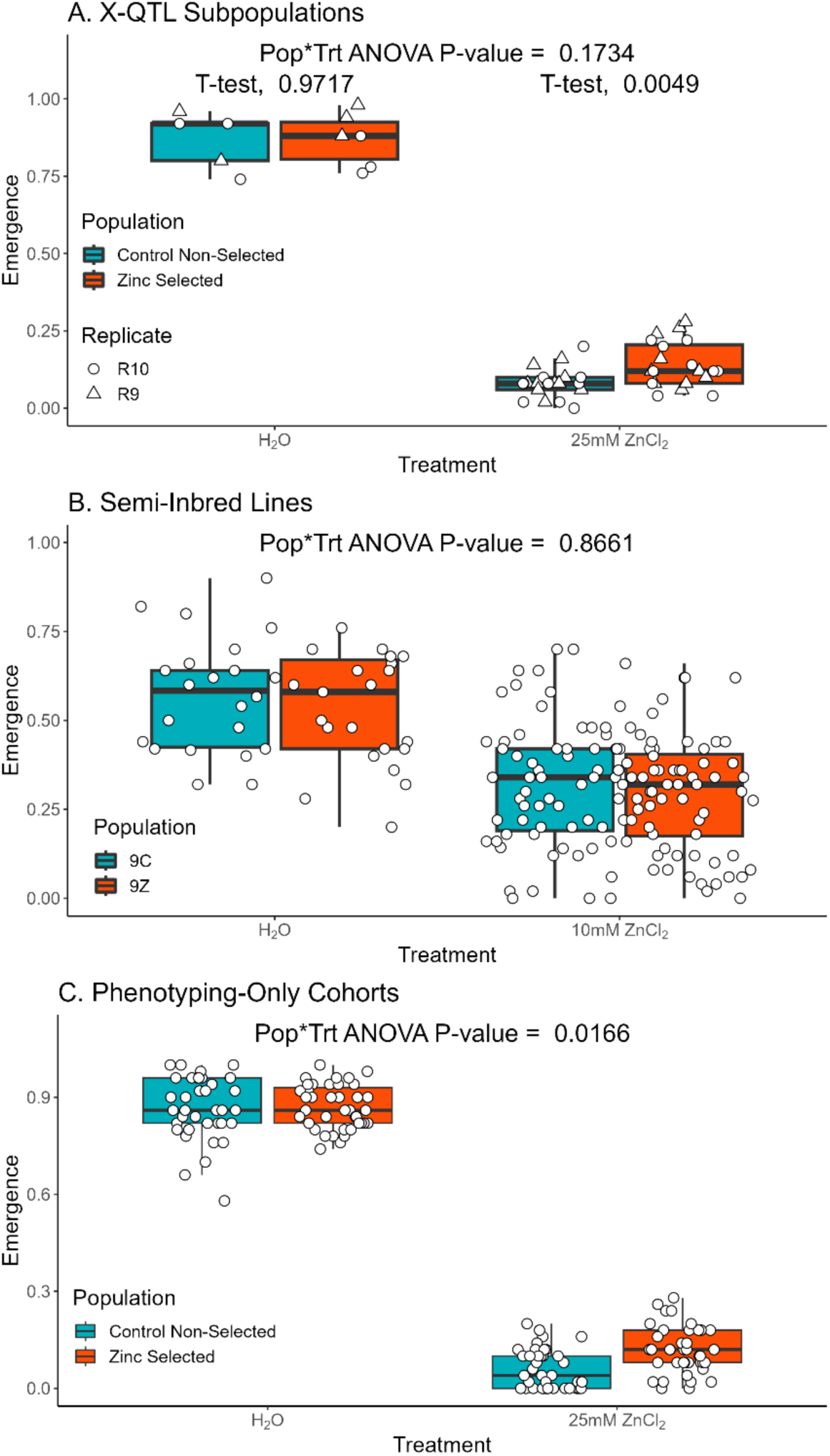
Zinc-selected populations show increased egg-to-adult emergence on zinc-containing media. We raised embryos from several different populations in both control media, and media containing a high zinc chloride concentration. Each point in the figure is the fraction of adults that emerged from a given test vial. (A) Outbred animals derived from X-QTL replicate 9 and 10 (R9, R10) samples; There is no significant population-by-treatment interaction, but zinc-selected populations show higher emergence on zinc media (*t*-test, *p* < 0.005). (B) Semi-inbred lines derived from the R9 and R10 Control and Zinc-selected cohorts; no significant population-by-treatment interaction. (C) Outbred animals generated in an analogous fashion to our X-QTL design; we see a significant population-by-treatment interaction (*p* < 0.02) with zinc-selected animals showing greater emergence on zinc-containing treatment. See Supplementary Table 3 for full details of the ANOVA statistical analyses.

**Figure 4:**
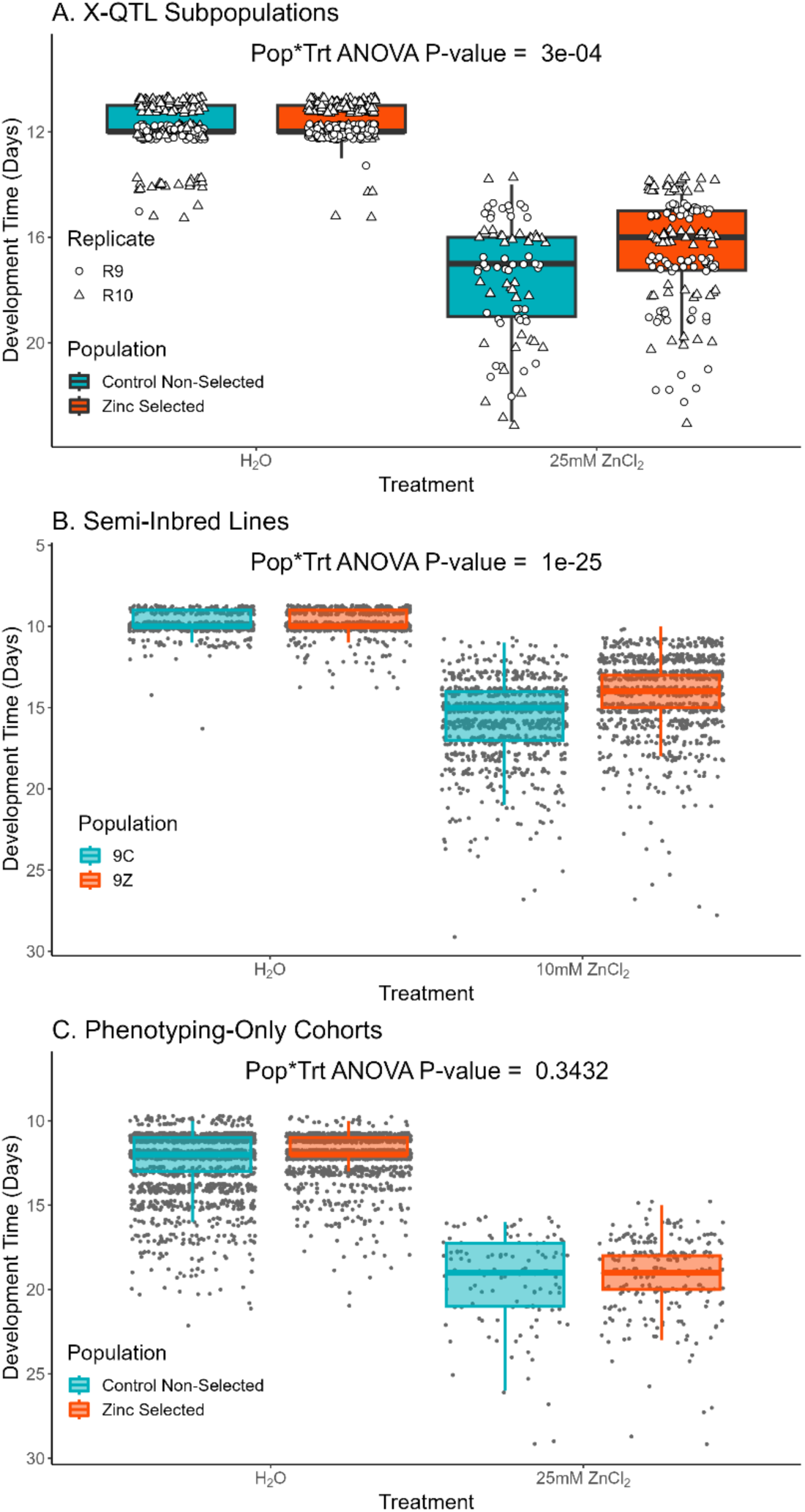
Zinc-selected populations show relatively faster development on zinc-containing media. We raised embryos from several different populations in both control media, and media containing a high zinc chloride concentration. Each point represents the egg-to-adult development time of a single animal. To more easily connect the development time data with emergence data in Figure 3 we have inverted the *y*-axes in these plots; points nearer the top of each panel reflect faster development. (A) Outbred animals derived from X-QTL replicate 9 and 10 (R9, R10) samples; Significant population-by-treatment interaction (*p* < 0.0005). (B) Semi-inbred lines derived from the R9 and R10 Control and Zinc-selected cohorts; Significant population-by-treatment interaction (*p* < 10^−25^). (C) Outbred animals generated in an analogous fashion to our X-QTL design; No significant population-by-treatment interaction. See Supplementary Table 4 for full details of the ANOVA statistical analyses.

### Evaluating the phenotypic impact of developmental zinc exposure

Exposure to high levels of ZnCl_2_ during the X-QTL selection assay resulted in fewer eggs surviving to adulthood (∼7% surviving) and an increase in egg-to-adult development time (of ∼5 days). To further explore the consequences of our zinc selection regime, we directly examined emergence (the percentage of eggs that develop into adults) and development time (the time, in days, between egg collection and adult emergence), in several different samples of animals derived from zinc selection and control, non-selection treatments: (1) Subpopulations of animals derived from the zinc-selected (Z), and the control, non-selected (C) animals that emerged in replicates R9 and R10 of the X-QTL mapping assay, (2) a series of partially (or semi-) inbred strains derived from the R9Z and R9C cohorts, and (3) animals derived via the same basic selection scheme used for the X-QTL mapping assay, but derived from a distinct base population established with just a fraction of the DSPR RILs, that yielded a pair of test cohorts we only employed for phenotyping. Our goal was to determine whether animals derived from selected and control groups responded differently to zinc treatment.

#### Developmental zinc exposure selects for genotypes with increased emergence on zinc media

Figure 3 shows the results of experiments examining egg-to-adult emergence on water control and zinc media for the 3 sets of populations described above (see Supplementary Table 3 for full ANOVA analysis results). For animals derived from our X-QTL selected and control, non-selected populations there was no population-by-treatment interaction (*p* = 0.17), potentially because this experiment had few water-control treatment vials (Figure 3A). However, animals from the zinc-selected cohorts did show increased emergence from zinc-containing media (*t*-test, *p* < 0.005, Figure 3A). The semi-inbred lines we constructed from our X-QTL cohorts showed no population-by-treatment interaction (*p* = 0.87, Figure 3B), suggesting that during the course of maintenance/inbreeding the subtle emergence phenotype may have been lost. Finally, the phenotyping-only test cohorts showed a significant population-by-treatment interaction (*p* = 0.02, Figure 3C). Overall, these results suggest that selection via developmental zinc exposure results in populations with increased egg-to-adult emergence on zinc-containing media.

#### Zinc-selected animals exhibit reduced developmental delay on zinc-containing media

Figure 4 presents the egg-to-adult development time results from the same experiments that led to Figure 3. Figure 4A highlights a significant population-by-treatment interaction (*p* < 0.0005, see Supplementary Table 4 for full ANOVA results); development time is similar for both X-QTL subpopulations in water control vials, but zinc-selected populations show shorter development time, i.e., exhibit a reduced developmental delay due to zinc. We see a similar result for the semi-inbred lines that were derived from the X-QTL subpopulations (Figure 4B, population-by-treatment interaction, *p* < 10^−25^), but see no effect in the phenotyping-only cohorts (Figure 4C), perhaps because relatively few animals emerged from zinc treatment vials in this experiment. Regardless, developmental selection for zinc resistance, in addition to enhancing emergence in zinc conditions (above and Figure 3), appears also to reduce the developmental delay that is a consequence of zinc treatment.

### Evaluating candidate genes via midgut-specific RNAi knockdowns

Of the 631 protein-coding genes within our X-QTL intervals, gene ontology information and data from previous fly studies of metal toxicity led us to 24 candidate genes (Supplementary Table 2). We were able to test the effect of 18 of these on developmental zinc resistance using a *mex1*-GAL4 driver which is expressed specifically in the midgut through all three stages of larval development (Binks et al., 2010; Phillips & Thomas, 2006; Tennessen et al., 2011). The midgut is arguably the relevant tissue for our target phenotypes since it is the primary site of nutrient absorption, and is where zinc absorption and homeostasis is primarily mediated (Lemaitre & Miguel-Aliaga, 2013; Navarro & Schneuwly, 2017).

RNAi knockdown of a gene can impact measured developmental phenotypes in the absence of any zinc challenge. Such events might indicate a gene influences the developmental response to zinc, but we are more interested in those genes that show a change in phenotype in response to a zinc challenge that *differs* between wildtype and knocked down gene expression levels. Analytically this is indicated by a significant interaction between genotype (UAS-RNAi knockdown *versus* wildtype genetic control) and treatment (zinc-containing *versus* water control treatments) and can be visualized by the slope of the lines connecting the mean phenotype on the two treatments for each genotype (see Figure 5).

**Figure 5:**
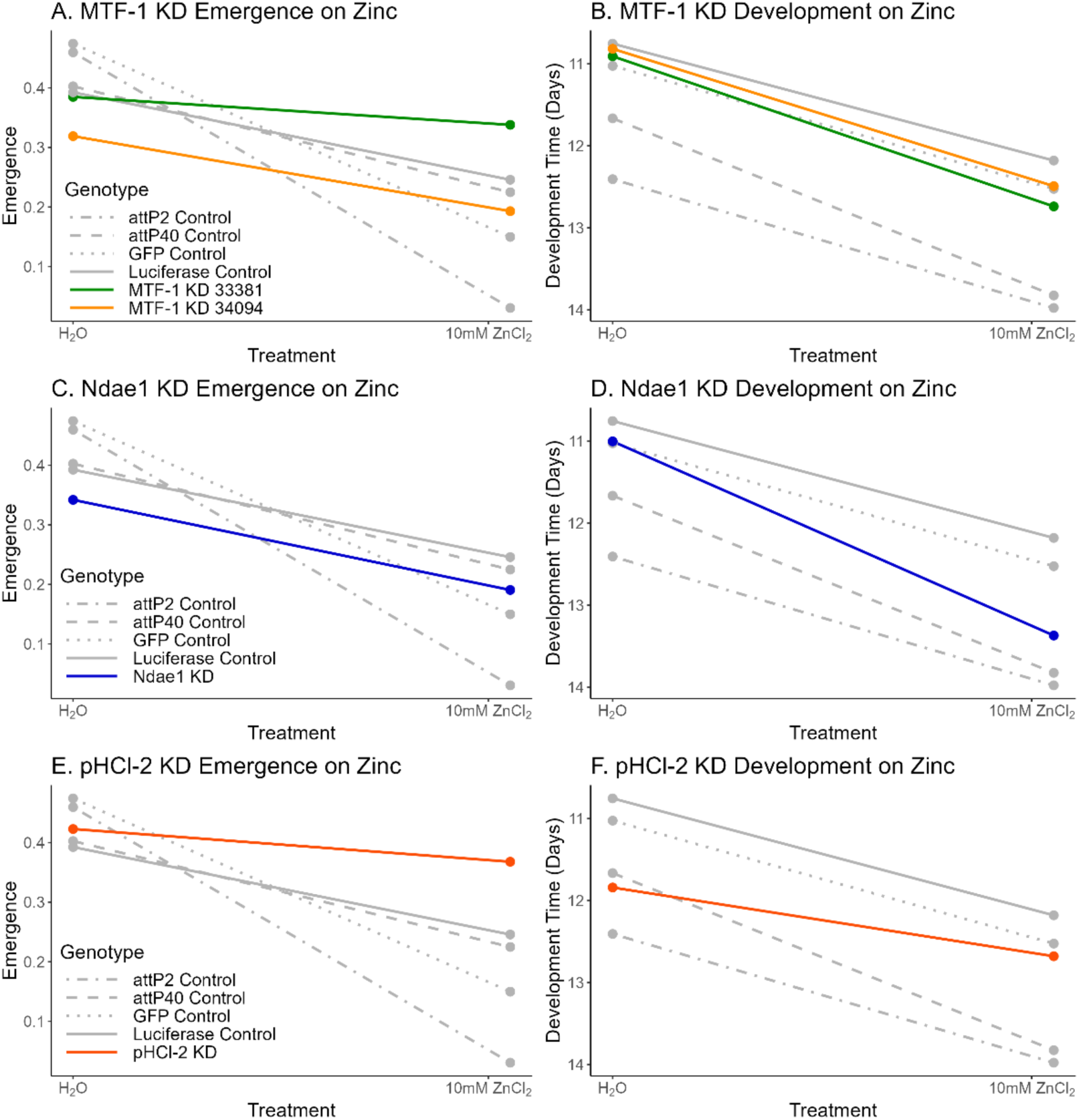
RNAi-based knockdowns of *MTF-1*, *Ndae1* and *pHCl-2* impact zinc resistance. Each genotype is the result of crossing a *mex1*-GAL4 strain with a strain that either carries a UAS-RNAi transgene (colored lines), or carries a control construct (gray lines). Each point represents the mean egg-to-adult emergence or development time for a given genotype for a given treatment. Confidence intervals around means were omitted for clarity, and to more easily focus on the differences in slopes between RNAi and control genotypes, which represent the genotype-by-treatment factor of interest (Supplementary Figure 4 does depict confidence intervals). Note the inverted *y*-axis used for the development time (right) panels; This was done such that a visibly negative slope for both emergence *and* development time yields the same inference; zinc treatment leads to a worse outcome (reduced emergence or longer development time.) *MTF-1* is an RNAi “hit” for emergence (panel A); knockdown using the 33381 transgene increases emergence relative to controls, as evidenced by a flatter slope. *Ndae1* is an RNAi “hit” for development time (panel D); knockdown increases developmental delay in zinc conditions as shown by a greater slope. *pHCl-2* is an RNAi “hit” for both emergence and development time (panels E and F); knockdown both increases emergence and reduces developmental delay, as evidenced by flatter slopes in both panels. See Table 2 and Supplementary Table 6 for statistical tests.

We employed multiple wildtype genetic controls to contrast with the UAS-RNAi knockdown genotypes; UAS-GFP and UAS-Luciferase strains express GFP or Luciferase, respectively, under UAS control, as well as strains that simply carry empty attP2 or attP40 docking sites (the positions where the UAS transgene is integrated). After crossing each of these to the *mex1*-GAL4 we saw significant variation in the emergence of progeny on zinc-supplemented media among the control genotypes, with the attP2-containing genotypes showing just 3% emergence on zinc-supplemented media *versus* 15-25% emergence for the other three control genotypes (Supplementary Figure 3, Supplementary Table 5). Given the wide variation among control genotypes, separately for each phenotype – emergence and development time – every RNAi knockdown genotype was contrasted with each of the four control genotypes in a separate full factorial ANOVA. In order to focus on the most interesting gene knockdown effects, RNAi “hits” for a given phenotype were called when the genotype-by-treatment interaction for at least one ANOVA was significant at *p* < 0.0003 (= 0.05/(21 × 4 × 2), since we tested 21 different UAS-RNAi genotypes, against 4 controls, for 2 phenotypes), the interaction was at least nominally significant (*p* < 0.05) for the other three ANOVAs, and the direction of the effect was consistent over the 4 tests.

We assessed the effect of midgut-specific RNAi-based knockdown for 18 genes using a set of 21 UAS-RNAi constructs (3 genes were tested with 2 different constructs). Ten genes emerged as hits. Results are summarized in Table 2 (see Supplementary Table 6 for full results of all analyses), and Figure 5 highlights results for 3 genes we discuss in more depth below (see Supplementary Figure 4 for similar plots for all genes tested). As shown in Table 2, 1 gene impacted only emergence, 7 impacted only development time, and 2 impacted both phenotypes (although for one of these genes – *Xrp1* – the effects on the two phenotypes were seen in different RNAi genotypes). The lack of consistency across the phenotypes implies the underlying genetics of the two traits may be at least partly distinct. Interestingly, our hits are enriched for those where expression knockdown appears to *increase* resistance. This is likely because some of our control, non-knockdown genotypes perform quite poorly on zinc media in our assay (Supplementary Figure 3), and it is therefore difficult for RNAi knockdowns to be statistically worse. We tested at least one candidate gene for each of the 7 mapped X-QTL (Table 1), and 5 QTL showed at least one hit (Table 2).Two of the genes with hits –*MTF-1* and *pHCl-2* – are associated with the “response to metal ion” GO term (GO:0010038), with *pHCl-2* specifically associated with the “cellular response to zinc ion” term (GO:0071294). Additionally, three of our hits – for *GluRIB*, *Ndae1*, and *Nup98-96* – were also hits for ZnCl_2_ response in a cell-based RNAi screen (Mohr et al., 2018). Notably, for *GluRIB* and *Nup98-96* our results matched observations from this cell-based screen in that knocking down expression of these genes *increases* resistance to zinc.

**Table 2.**
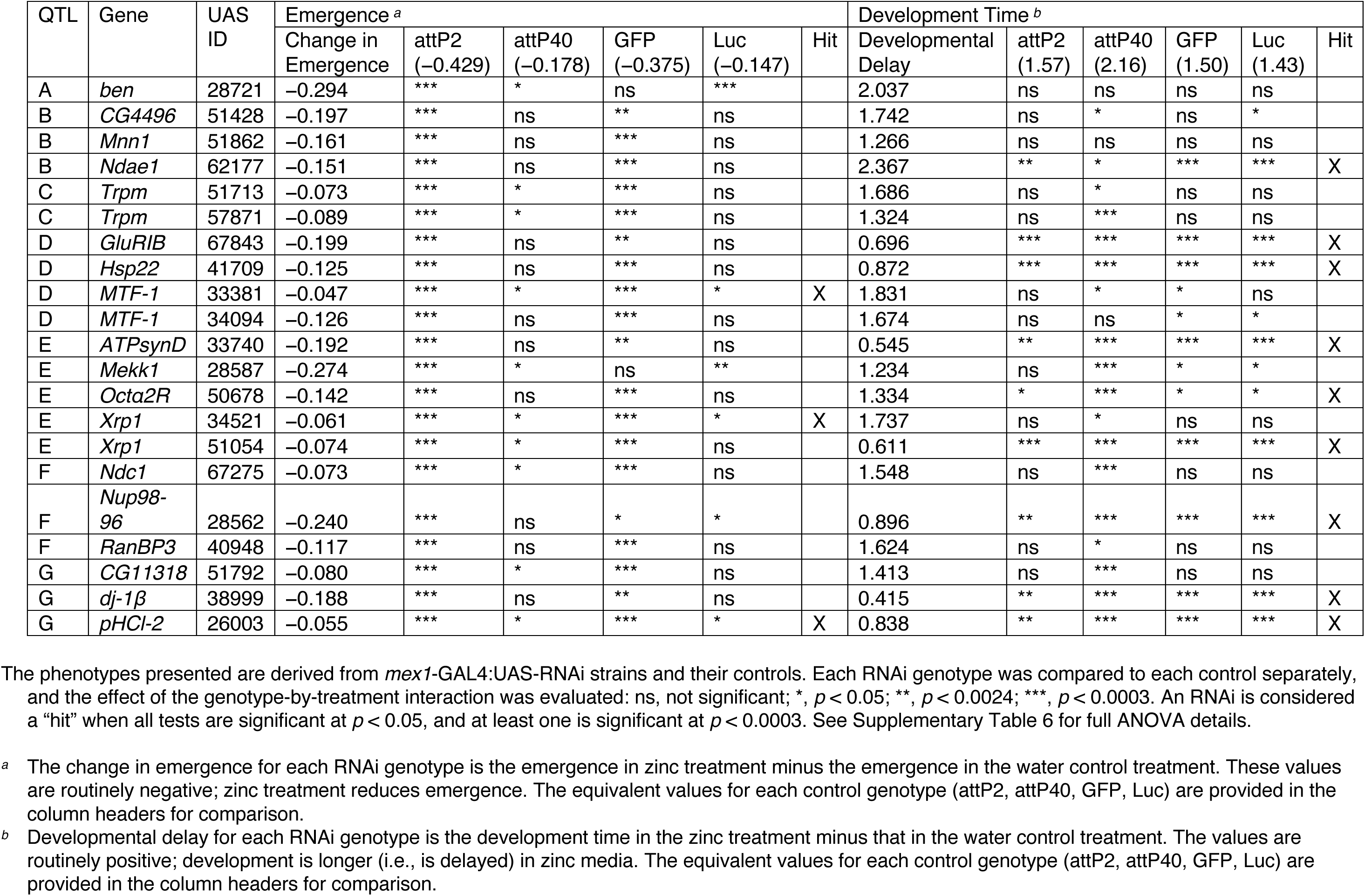
Effects of RNAi knockdown on emergence and development time.

### Knockdown of *MTF-1* in the midgut increases egg-to-adult emergence on zinc media

*MTF-1* encodes a metal transcription factor that, in addition to playing a role in the detoxification of a range of metals, is involved in zinc homeostasis and detoxification, and induces the expression of zinc transporters and metallothionein proteins (Balamurugan et al., 2004; Egli et al., 2003; Navarro & Schneuwly, 2017). We knocked down expression of *MTF-1* using two different UAS-RNAi constructs, and found a significant genotype-by-treatment emergence effect for UAS ID 33381 (Table 2, green line in Figure 5A); compared to all four genetic controls, this *MTF-1* knockdown shows enhanced emergence in media supplemented with zinc. The second UAS (ID 34094) shows the same trend, but the genotype-by-treatment interaction is not significant in all cases (Table 2). There was no evidence for an effect of either *MTF-1* knockdown on development time (Table 2, Figure 5B).

That knocking down the expression of *MTF-1* results in increased resistance in our study is somewhat counterintuitive since a number of studies have shown that *MTF-1* loss-of-function mutations result in susceptibility to the toxic effects of metals (Bahadorani et al., 2010; Balamurugan et al., 2004; Egli et al., 2003; Wang et al., 2004). This being said, Bahadorani et al. (Bahadorani et al. 2010) showed that while *MTF-1* null flies were more susceptible to cadmium, iron and copper toxicity, nulls showed no difference in zinc susceptibility compared to wildtype. The same study also showed that overexpression of *MTF-1* made flies more susceptible to zinc exposure both as adults (flies died more quickly) and as larvae (fewer eggs developed into adults), in contrast to the result for other metals, where *MTF-1* overexpression increased resistance. Combined with our results, this work suggests there may be a more complex relationship between zinc toxicity resistance and *MTF-1* expression level.

### Reduced *Ndae1* expression results in greater developmental delay in zinc media

*Ndae1* encodes an Na^+^-driven anion exchange membrane protein found in the nervous system, throughout the digestive tract, and in the malpighian tubules (Romero et al., 2000; Sciortino et al., 2001). In a RNAi screen using *Drosophila* cultured cells exposed to toxic zinc conditions, *Ndae1* was identified as a high confidence hit whose knockdown led to increased zinc resistance as measured by ATP level, an indirect readout of cell viability (Mohr et al., 2018).

In our tests, the impact of *Ndae1* knockdown on emergence was dependent on the genetic control to which it is compared (Table 2, Figure 5C). However, *Ndae1* knockdowns showed significant genotype-by-treatment effects for development time against all 4 control genotypes (Table 2, Figure 5D), with reduced expression resulting in greater development time in zinc media, i.e., increased susceptibility (see the greater slope of the blue *Ndae1* RNAi line in Figure 5D). This reduction in resistance appears to contradict the observations made by Mohr et al (Mohr et al., 2018). This discrepancy could be the use of different models (whole animals *versus* cultured cells), or different readouts (direct measures of emergence and development time *versus* a proxy for cell viability), and would require additional investigation to understand.

### RNAi-based *pHCl-2* expression knockdown results in lower emergence and greater development time in zinc media

*pHCl-2* encodes a zinc-gated chloride channel that is expressed in the midgut, concentrated in the copper cell region (an acid-secreting region of the midgut, sometimes considered the fly “stomach”), and in the malpighian tubules (Feingold et al., 2016; Redhai et al., 2020; Remnant et al., 2016). Previous work suggests that *pHCl-2* is a zinc sensor and that it may play a role in directing larvae to nutrient-rich food (Redhai et al., 2020).

*pHCl-2* impacted emergence on zinc media, with the *pHCl-2* midgut-specific knockdown genotype showing greater emergence than controls in zinc conditions (Table 2, Figure 5E), i.e., *pHCl-2* knockdown appears to increase resistance to the toxic effects of zinc. These results mirror data from similar experiments testing *pHCl-2* (known previously as CG11340) gut RNAi knockdown and gene knockouts on media supplemented with copper; *pHCl-2* knockdown/knockout increases survival compared to the control (Remnant et al., 2016). We additionally found significant genotype-by-treatment interactions for development time for all *pHCl-2* RNAi versus genetic control contrasts (Table 2), with the knockdown genotype showing reduced developmental delay in response to zinc media (see lower slope of red line in Figure 5F), again indicating lower expression of *pHCl-2* yields zinc resistance.

## DISCUSSION

In this study, we employed an “extreme” or X-QTL mapping approach to identify regions of the genome harboring variants contributing to developmental zinc resistance in *Drosophila melanogaster*. We successfully mapped several such QTL across the major *D. melanogaster* chromosome arms, with five of the QTL - all present in euchromatic genomic regions having normal levels of recombination - resolving to relatively modest genomic intervals of 300-820kb, or 0.65-2.85cM (see Table 1). The mapping power and resolution we achieved in the present study is higher than in our previous caffeine resistance X-QTL work (Macdonald et al., 2022), which we attribute both to the greater number of experimental replicates employed (12 versus 4) and the larger number of individuals in the DNA pools (an average of 302 versus an average of 245). This highlights the value of increased replication and pool size in X-QTL designs for improving QTL detection and localization, as we previously claimed via simulations (Macdonald et al., 2022).

### Zinc toxicity resistance represents a multi-faceted phenotype

Our study suggests that zinc toxicity resistance is a complex trait encompassing multiple components, primarily egg-to-adult viability and development time. However, the nature of resistance appears to extend beyond these two aspects. Through our experiments with different populations/strains, we observed that the components of zinc resistance can be dissociated. For instance, semi-inbred lines derived from zinc-selected populations maintained resistance in terms of developmental time but lost the emergence rate advantage (compare Figures 3B and 4B), possibly due to genetic drift occurring during establishment of the lines. Conversely, the zinc-selected, phenotyping-only cohort showed improved emergence rates on zinc media but no difference in developmental time (compare Figures 3C and 4C). These observations suggest that developmental time and emergence rate under zinc stress are independent phenotypes, a notion further supported by our RNAi results where gene knockdowns could affect either or both phenotypes (see Table 2).

There also appears to be a distinction between zinc resistance during development and that during adulthood; Adult females from populations that experienced developmental zinc selection showed no increased resistance to zinc, and in some cases even exhibited a slightly higher mortality rate during zinc exposure (Supplementary Figure 5). This underscores the importance of considering life stage-specific responses when studying metal toxicity (Everman et al., 2021). The complexity of zinc resistance is further highlighted by the potential concentration-dependent effects we observed. Our X-QTL mapping used 25mM ZnCl_2_, while subsequent phenotyping of semi-inbred lines and RNAi knockdowns necessitated a lower concentration of 10mM ZnCl_2_ due to the extreme toxicity of the higher dose for those. The difference in concentration could contribute to variation in observed phenotypes, and re-emphasizes the importance of considering dose-dependent responses in toxicity studies (Knowles et al., 2018; Wang & Kruglyak 2014; Widmayer et al., 2022). Collectively, our findings suggest zinc toxicity is a complex trait with multiple, potentially independent components that can vary across developmental stages and zinc concentrations.

### Impact of selection for zinc resistance on susceptibility to other heavy metals

The zinc resistance loci we identify here do not overlap in location with those previously mapped for caffeine and malathion resistance using the same base population (Macdonald et al., 2022; Macdonald & Long, 2022). With the caveat that we do not have perfect power in any of these studies, this suggests that the genetic basis of zinc resistance is distinct from these other xenobiotic toxicity traits. However, one of the zinc resistance QTL (QTL-E) does colocalize with a locus contributing to copper resistance identified in an independent set of DSPR RILs (Everman et al., 2021), potentially hinting that there are some shared mechanisms of resistance between zinc and copper, two heavy metals, as has been previously suggested in *C. elegans* (Evans et al., 2020).

To explore the specificity of the zinc resistance in our populations, we tested our semi-inbred lines and phenotyping-only cohorts against toxic concentrations of copper (2mM CuSO_4_) and cadmium (0.2mM CdCl_2_) in addition to a zinc challenge. In the semi-inbred lines raised on zinc media, those lines derived from the zinc-selected population exhibit a reduced developmental delay (Figure 4B), but this pattern is not recapitulated when they are raised on copper or cadmium media (Supplementary Figure 7B). In the phenotyping-only cohorts, when raised on zinc media the zinc-selected cohort has a higher rate of emergence than the control cohort (Figure 3C), whereas when raised on cadmium medium the opposite pattern is observed; zinc selection reduces emergence on cadmium media (Supplementary Figure 6A). Additionally, while the zinc-selected and control phenotyping-only cohorts do not differ in development time on zinc media (Figure 4C), the zinc-selected cohorts show slower development on cadmium and copper media than the control cohorts (Supplementary Figure 7A). These results suggest that in some instances the genetic mechanisms underlying zinc resistance may confer susceptibility to other heavy metals.

Contrasting effects in resistance between zinc and other heavy metals have been documented previously. For example, Bahadorani et al. (2010) found that MTF-1 overexpression had the opposite effect on zinc resistance compared to other heavy metals like cadmium and copper. Such differences may be related to distinct chemical properties of the metals. Zinc is redox neutral in biological systems, unlike copper and iron, and only promotes formation of reactive oxygen species at high, toxic levels (Harold, 2014; Kambe et al., 2015; Maret, 2013; Navarro & Schneuwly, 2017). Metals such as copper may induce toxicity response pathways that our zinc-selected flies may not be adapted to handle effectively. In terms of cadmium, studies in *C. elegans* suggest that cadmium toxicity can cause zinc deficiency by activating zinc detoxification pathways (Earley et al., 2021; Moyson et al., 2018; Power & de Pomerai, 1999). Earley et al. (2021) demonstrated that the major pathways governing zinc toxicity resistance are not significantly involved in cadmium resistance. This separation of pathways could explain why our zinc-selected flies are not better equipped to survive cadmium exposure, and may even be more susceptible to zinc deficiency induced by cadmium. Additional studies using our X-QTL design seeking to characterize the genetic basis of variation in susceptibility to a range of metals would enable better examination of metal-specific genetic adaptations.

### Mapped X-QTL do not routinely harbor known zinc toxicity/homeostasis genes

Our X-QTL mapping revealed seven loci contributing to zinc toxicity resistance, with only three intervals containing genes previously associated with zinc homeostasis: *Trpm* in QTL-C, *MTF-1* in QTL-D, and *MtnF* in QTL-F. Perhaps surprisingly, most known zinc transporters were not found within our QTL, despite them commonly being focus of zinc toxicity and homeostasis research. One potential explanation is that there is limited functionally-relevant standing variation at such genes, making them harder to detect via our mapping design. Regardless, the observation highlights the potential importance of other gene families in mediating zinc resistance, and underscores the value of unbiased, genome-wide approaches like X-QTL mapping in uncovering novel factors contributing to metal toxicity resistance. Furthermore, by delivering QTL intervals that implicate only a modest number of genes, X-QTL can enable subsequent characterization of novel causative loci.

### Midgut-specific RNAi knockdowns identify genes important in zinc toxicity, while challenging some extant assumptions

Our RNAi-based functional testing of candidate genes within the QTL intervals yielded several interesting results. We identified ten genes that, when knocked down in the midgut, significantly affected egg-to-adult emergence, development time, or both traits, in zinc-supplemented media. Perhaps surprisingly, for all but one of our hits, knocking down the gene resulted in greater (rather than lower) resistance to zinc. We speculate that the concentration of zinc chloride we chose to apply in these assays was sufficiently high that it was unlikely for a knockdown genotype to perform significantly worse than the control genotype; A similar effect was noted in a cell-based RNAi screen for zinc toxicity genes, where they suspected the zinc treatment applied was so toxic that it was difficult for worse, RNAi-generated phenotypes to be detected (Mohr et al., 2018).

Some of the genes identified, such as *GluRIB* and *Nup98-96*, showed effects consistent with a previous cell-based RNAi screen for zinc toxicity (Mohr et al., 2018), providing cross-validation of their role in zinc resistance. We contend that our decision to employ an RNAi knockdown strategy that is specific to the midget (Phillips & Thomas, 2006) makes our conclusion that these genes are involved in zinc resistance more robust than had we used a less tissue specific driver (e.g., a ubiquitous actin- or tubulin-based RNAi knockdown), where unintentional off-target effects could be more likely to bias results. This being said, RNAi knockdowns do not prove that standing variation at the genes we identify drive the initial X-QTL mapping result. Allele replacements via CRISPR/Cas9 would provide more concrete evidence, but are considerably more difficult to achieve.

The behavior of MTF-1 knockdowns in our RNAi screen was particularly intriguing. Contrary to expectations based on its known role in metal detoxification, in our study knockdown of MTF-1 in the midgut led to increased emergence on zinc-supplemented media. This result, combined with previous findings showing increased zinc susceptibility with MTF-1 overexpression (Bahadorani et al., 2010), suggests a more complex relationship between MTF-1 expression levels and zinc resistance than previously thought. Further investigation into the dose-dependent effects of MTF-1 on zinc homeostasis and resistance could yield valuable insights into the regulation of metal detoxification pathways.

### Founder haplotypes showing the greatest changes in frequency in zinc-selected samples tend to be common in the base population

The observation that founder haplotypes showing substantial changes in frequency due to zinc selection are consistently those at high frequency in the base population is novel. This pattern could reflect the actions of random genetic drift and selection during the creation and maintenance of the DSPR RILs (which we mix to yield our X-QTL base population), or it could indicate that resistance/susceptibility alleles are relatively common in natural populations. In contrast, this observation may have been predictable on theoretical grounds. The change in a founder haplotype frequency in the selected pool versus the base population is proportional to *pqs*, where *p* is the frequency of the haplotype in the base population, *q* is 1 – *p*, and *s* is the strength of selection on the variant (i.e., its effect size). Thus, for a given effect size, we expect more common alleles to show bigger allele frequency shifts, and a corollary is that if the founder allele frequencies in the base population are not each close to 1/*F* (where *F* is the number of founders) the number of alleles effectively queried is less than *F*. Further investigation into the frequencies of these alleles in diverse natural populations will be required to provide insights into the evolutionary history and adaptive significance of zinc resistance in *Drosophila*.

## Conclusion

Our X-QTL mapping approach, combined with a targeted, tissue-specific RNAi screen, has revealed several genomic regions and specific genes contributing to variation in developmental zinc resistance in *Drosophila*. These findings not only advance our understanding of the genetic architecture of metal toxicity resistance, but also highlight the complex and sometimes counterintuitive nature of metal homeostasis mechanisms. Future studies exploring the molecular mechanisms of these identified genes, their potential roles in resistance to other metals, and their evolutionary significance in natural populations will further elucidate the intricate processes underlying metal toxicity resistance in insects and potentially other organisms.

## ACKNOWLEDGEMENTS

This work was supported by NIH awards R01-ES029922 (to SJM) and R01-OD034064 (to SJM and ADL). Computational infrastructure and assistance to authors at KU was provided by the Data Science Core of the Kansas INBRE project (supported by NIH P20-GM103418). The authors acknowledge the Bloomington *Drosophila* Stock Center (P40-OD018537) for supplying strains used in the project.

