## Supplementary Materials for "High-throughput genetic mapping discovers novel zinc toxicity response loci in *Drosophila melanogaster*"

**Supplementary Table 1:** Details of all 12 X-QTL replicates.

| Replicate | Non-Selection Control Treatment | | | | Zinc Selection Treatment | | | | Estimated Number of Females Exposed to Zinc Treatment * | Estimated Fraction of Females Exposed to Zinc and Survived (%) * |
|---|---|---|---|---|---|---|---|---|---|---|
| | Number of Bottles | Embryos/ Bottle (µl) | Total Emerged Females per Replicate | Emerged Females per µl | Number of Bottles | Embryos/ Bottle (µl) | Total Emerged Females per Replicate | Emerged Females per µl | | |
| R1 | 4 | 24 | 523 | 5.448 | 20 | 48 | 376 | 0.392 | 5230 | 7.189 |
| R2 | 4 | 24 | 498 | 5.188 | 17 | 48 | 329 | 0.403 | 4233 | 7.772 |
| R3 | 4 | 24 | 401 | 4.177 | 18 | 48 | 299 | 0.346 | 3609 | 8.285 |
| R4 | 4 | 24 | 479 | 4.990 | 21 | 48 | 260 | 0.258 | 5029.5 | 5.169 |
| R5 | 4 | 24 | 437 | 4.552 | 12 | 48 | 192 | 0.333 | 2622 | 7.323 |
| R6 | 4 | 24 | 486 | 5.063 | 20 | 48 | 190 | 0.198 | 4860 | 3.909 |
| R7 | 4 | 24 | 600 | 6.250 | 20 | 48 | 476 | 0.496 | 6000 | 7.933 |
| R8 | 4 | 48 | 1017 | 5.297 | 12 | 60 | 351 | 0.488 | 3813.75 | 9.204 |
| R9 | 4 | 24 | 754 | 7.854 | 8 | 48 | 169 | 0.440 | 3016 | 5.603 |
| R10 | 4 | 24 | 725 | 7.552 | 12 | 48 | 290 | 0.503 | 4350 | 6.667 |
| R11 | 4 | 24 | 638 | 6.646 | 12 | 48 | 311 | 0.540 | 3828 | 8.124 |
| R12 | 4 | 24 | 739 | 7.698 | 14 | 48 | 379 | 0.564 | 5173 | 7.327 |

* To enhance efficiency, we did not manually count the number of embryos tested per bottle/replicate/treatment, and instead pipetted aliquots of embryos resuspended in 1X PBS into each bottle. Given we do not know the number of embryos per bottle, we are estimating the zinc-selection pressure exerted on females based on the emergence of control females within each replicate. For each treatment the “Emerged Females per µl” values are the “Total Number of Emerged Females per Replicate” divided by the total volume of embryos (in µl) used per replicate (“Number Bottles” multiplied by “Embryos/Bottle (µl)”). The “Estimated Number of Females Exposed to Zinc Treatment” is the total volume of embryos (in µl) used for the Zinc Selection Treatment multiplied by the number of “Emerged Females per µl” for the Non-Selection Control Treatment. And the “Estimated Fraction of Females Exposed to Zinc and Survived (%)” is calculated as the ratio of the “Emerged Females per µl” for the two treatments (Zinc Selection divided by Non-Selection Control).

**Supplementary Text 1:** DNA isolation protocol.

Requires Qiagen Puregene Cell Kit (Cat. No. 158046) for the Cell Lysis Solution, RNase A, and Protein Precipitation Solution, and Qiagen Buffer EB (Cat. No. 19086).

1. Add 5 glass beads and 1ml of 1X PBS to a 2ml screwtop tube containing the pool of flies. Then homogenize the flies by placing tube in a MiniBeadBeater-96, running the instrument for two 45 second intervals.

2. Pour homogenate into a 15ml tube, then rinse out the original screwtop tube with 1ml of cold Cell Lysis Solution (CLS) and then pour into the 15ml tube. Repeat as necessary.

3. Add 7 ml of CLS, minus the amount used to rinse the screwtop tube (see Step 2), to the 15 ml tube. Invert to mix, and then incubate at 65°C in a hybridization oven for 30 minutes.

4. Cool to room temperature and then move 600 µl of homogenate to 1.7ml tube. (The remainder of the homogenate can be discarded, or frozen at −20°C in case additional DNA is needed subsequently).

5. Add 3 µl of undiluted RNase A solution to the 600 µl of homogenate, mix, and then incubate in a hot block at 37°C for 40 minutes. Rapidly cool sample to room temperature by placing on ice for 1 minute.

6. Add 200 µl protein precipitation solution to the sample, vortex on high speed for 20 seconds, and place sample on ice for 5 minutes. Centrifuge sample at ~20,000 g for 3 minutes.

7. Move 600 µl of the supernatant to a new 1.7 ml tube containing 600 µl of Isopropanol and mix by inverting. Centrifuge at ~20,000 g for 1 minute, and gently pour off supernatant.

8. Add 600 µl of 70% ethanol, and invert tube to wash pellet. Centrifuge at ~20,000 g, for 1 minute, and gently pour off supernatant. Invert tube on absorbent paper and air dry for 15 minutes.

9. Add 50 µl EB buffer and incubate at 65 °C for 1 hour. Flick the tube to resuspend DNA.

**Supplementary Table 2:** Candidate genes in X-QTL intervals.

| X-QTL | Gene Symbol | Gene Name | FlyBase ID |
|---|---|---|---|
| A | ben | bendless | FBgn0000173 |
| B | Gart ** | GART trifunctional enzyme | FBgn0000053 |
| B | Mnn1 | Menin 1 | FBgn0031885 |
| B | CG4496 | - | FBgn0031894 |
| B | Ndae1 | $Na^{+}$-driven anion exchanger 1 | FBgn0259111 |
| C | Trpm | Transient receptor potential cation channel, subfamily M | FBgn0265194 |
| C | Vha36-1 ** | Vacuolar H+ ATPase 36kD subunit 1 | FBgn0022097 |
| C | Vha14-1 ** | Vaculoar H+ ATPase 14kD subunit 1 | FBgn0262512 |
| D | GluRIB | Glutamate receptor IB | FBgn0264000 |
| D | Hsp22 | Heat shock protein 22 | FBgn0001223 |
| D | MTF-1 | Metal response element-binding Transcription Factor-1 | FBgn0040305 |
| E | CG14302 ** | - | FBgn0038647 |
| E | Mekk1 | Mekk1 | FBgn0024329 |
| E | Octα2R | α2-adrenergic-like-octopamine receptor | FBgn0038653 |
| E | Xrp1 | Xrp1 | FBgn0261113 |
| E * | ATPsynD | ATP synthase, subunit D | FBgn0016120 |
| F | RanBP3 | Ran binding protein 3 | FBgn0039110 |
| F | Nup98-96 | Nucleoporin 98-96kD | FBgn0039120 |
| F | Ndc1 | Nuclear division cycle 1 | FBgn0039125 |
| G | Fer1HCH *** | Ferritin 1 heavy chain homologue | FBgn0015222 |
| G | Fer2LCH *** | Ferritin 2 light chain homologue | FBgn0015221 |
| G | dj-1β | dj-1β | FBgn0039802 |
| G * | pHCl-2 | pH-sensitive chloride channel 2 | FBgn0039840 |
| G * | CG11318 | - | FBgn0039818 |

Information sourced from FB2022_06 (Gramates et al. 2022).

* These 3 genes are each just outside the formal 3-LOD support interval for our final set of X-QTL (see Table 1). These genes are included in our candidate set because they were within X-QTL intervals using an earlier version of our X-QTL analysis framework, and remain very close to the final set of X-QTL intervals we report.

** No TRiP UAS-RNAi strains were available for these target genes.

*** Knockdown of these genes via *mex1*-GAL4-driven, midgut-specific RNAi led to almost zero adult emergence in the water control treatment in initial tests, suggesting gene knockdown is lethal. These genes were not assayed further.

**Supplementary Text 2.** Ingredients and brief protocol for Macdonald lab cornmeal-yeast-molasses fly media.

28.5-liters water
318-g agar (Genesee Scientific; 66-111. This is based on a gel strength of 960 g/cm2, and will change depending on the batch)

- Add water to steam kettle, turn on electric mixer, and slowly add agar
- Bring mix to a boil

3,200-ml molasses (Genesee Scientific; 62-117)

- Reduce the kettle pressure to reduce the heat slightly
- Add molasses, and bring mix back to a boil

4-liters water
1,460-g inactive dry yeast (Genesee Scientific; 62-107)

- Mix in bucket using paint-stirring drill attachment

4-liters water
2,600-g yellow cornmeal (Genesee Scientific; 62-101)

- Mix in bucket using paint-stirring drill attachment
- Add both the water/yeast and water/cornmeal mixes to the steam kettle
- Bring mix back to boil, and simmer for ~15-min
- Release pressure from steam kettle, but continue to stir with electric mixer

330-ml water
259-ml propionic acid (ThermoFisher; A258-500)
31-ml phosphoric acid (85%; ThermoFisher; A242-500)

- Pour mix into kettle

400-ml 95% ethanol
40-g tegosept (Genesee Scientific; 20-258)

- Dissolve tegosept in ethanol
- Pour mix into kettle
- Fill vials/bottles

**Supplementary Figure 1:** Comparing duplicate control pools reveals few differences.

For five X-QTL replicates – R5, R6, R8, R11 and R12 – we generated sequencing libraries from two independent pools of unselected, control females ("A" and "B" pools for each replicate). We tested for differentiation in the haplotype frequencies between the pairs of control pools following the same analysis used to compare zinc-selected and unselected populations, treating the "A" and "B" sets of pools as different treatments. No peaks reached above a $-\log_{10}(P)$ threshold of 4. On the X chromosome there is one aberrant window with a $-\log_{10}(P)$ value >5, but this point does not fall within the interval implicated by QTL-A (the only X-linked zinc resistance locus we identified). Equally, the peak on chromosome 3L that has a $-\log_{10}(P)$ value of ~3.5 does not full within the LOD drop interval for QTL-D.

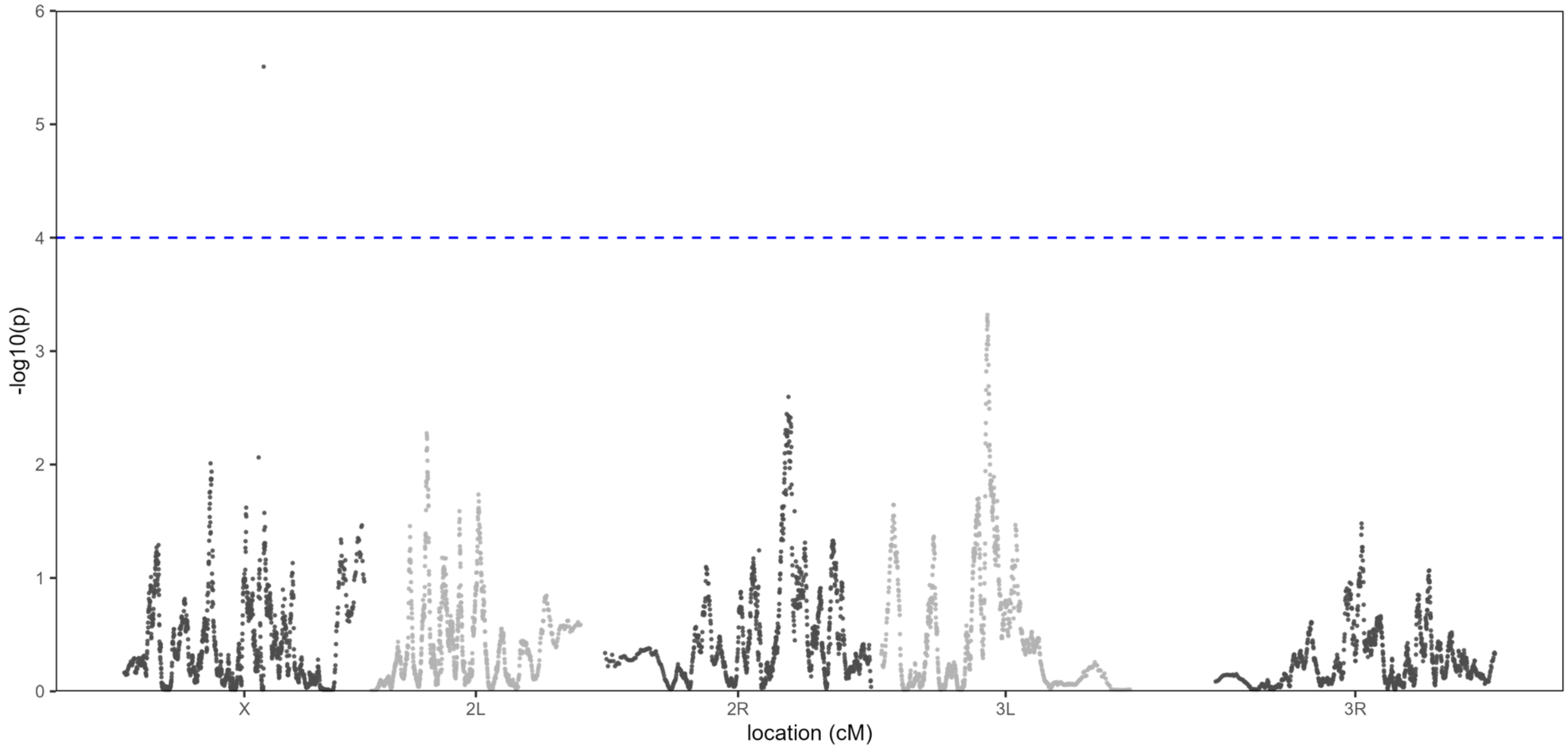

**Supplementary Figure 2:** Founder haplotype frequencies at mapped X-QTL.

Each of the 7 multi-panel figures on the next few pages depicts a single QTL (A-G) and comprises 4 panels. The top panel is identical to the relevant panel from Figure 2 in the main text, and shows the haplotype frequency difference (selection minus control) centered at each QTL. The middle panels show the estimated frequency for each founder haplotype separately for the control (left) and selected (right) populations through the QTL interval. The bottom panel shows the frequencies for each founder in the selected and control populations directly at the QTL peak. For QTL-A, the top panel shows the QTL is driven by A1 (carrying a resistance allele) and A7 (carrying a susceptibility allele). The remaining 6 founders show no/limited frequency change. The middle/bottom panels show that A7 and A1 are among the most common founders at this location in the genome. The remaining figures highlight a similar pattern for the other mapped QTL; QTL are driven by the more common founders at those genomic locations.

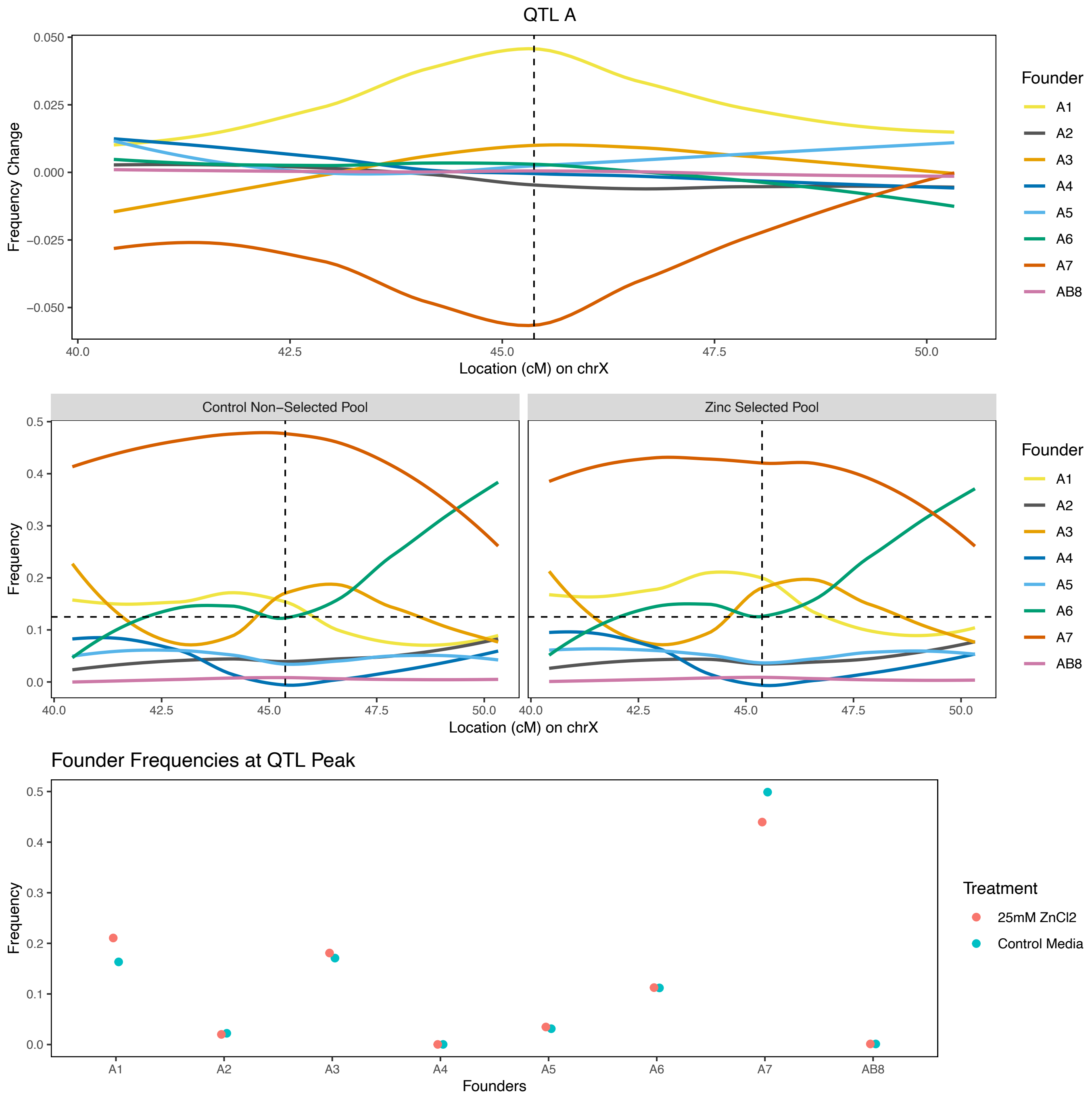

**Supplementary Figure 2:** Contd.

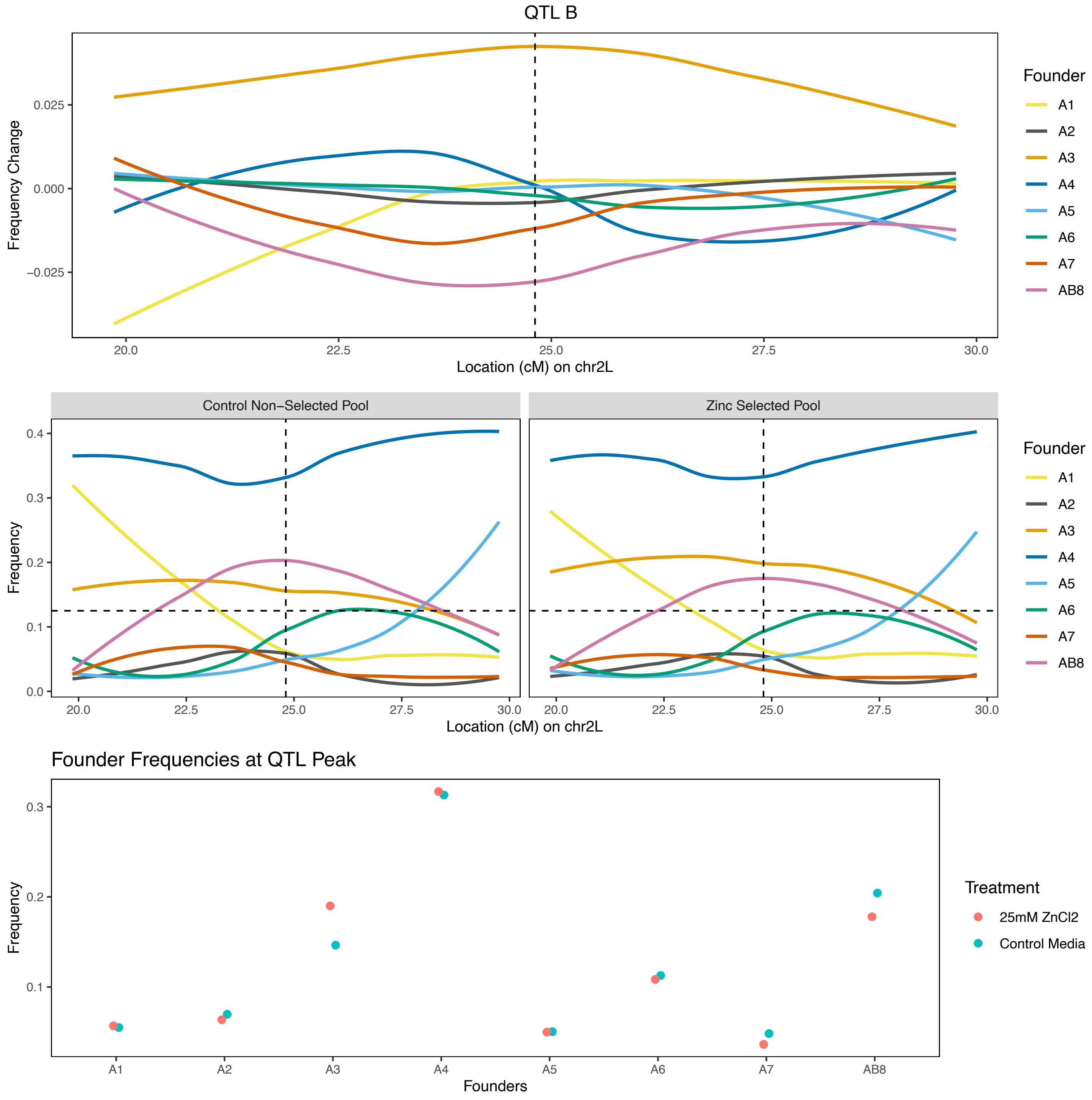

**Supplementary Figure 2:** Contd.

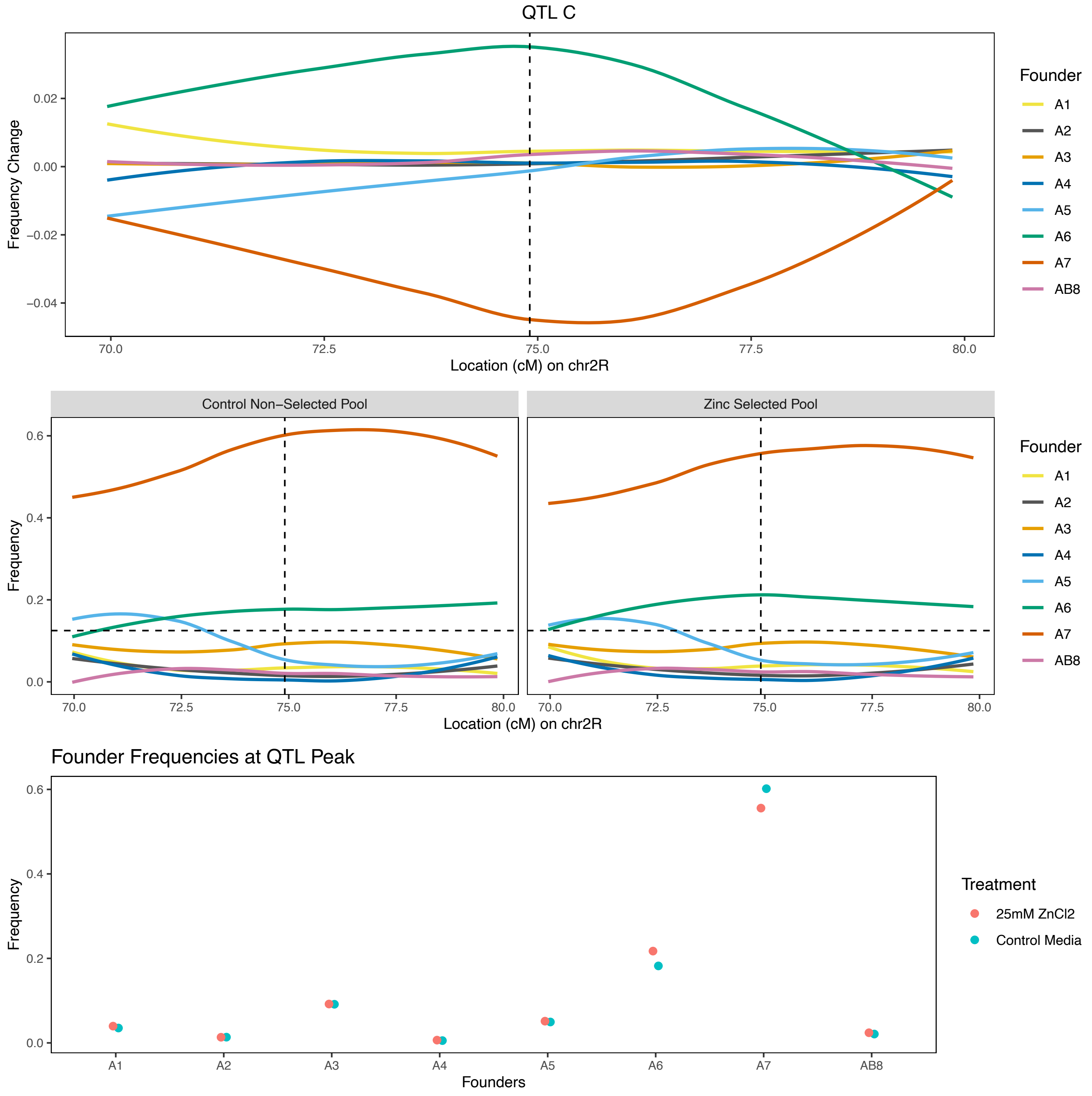

**Supplementary Figure 2:** Contd.

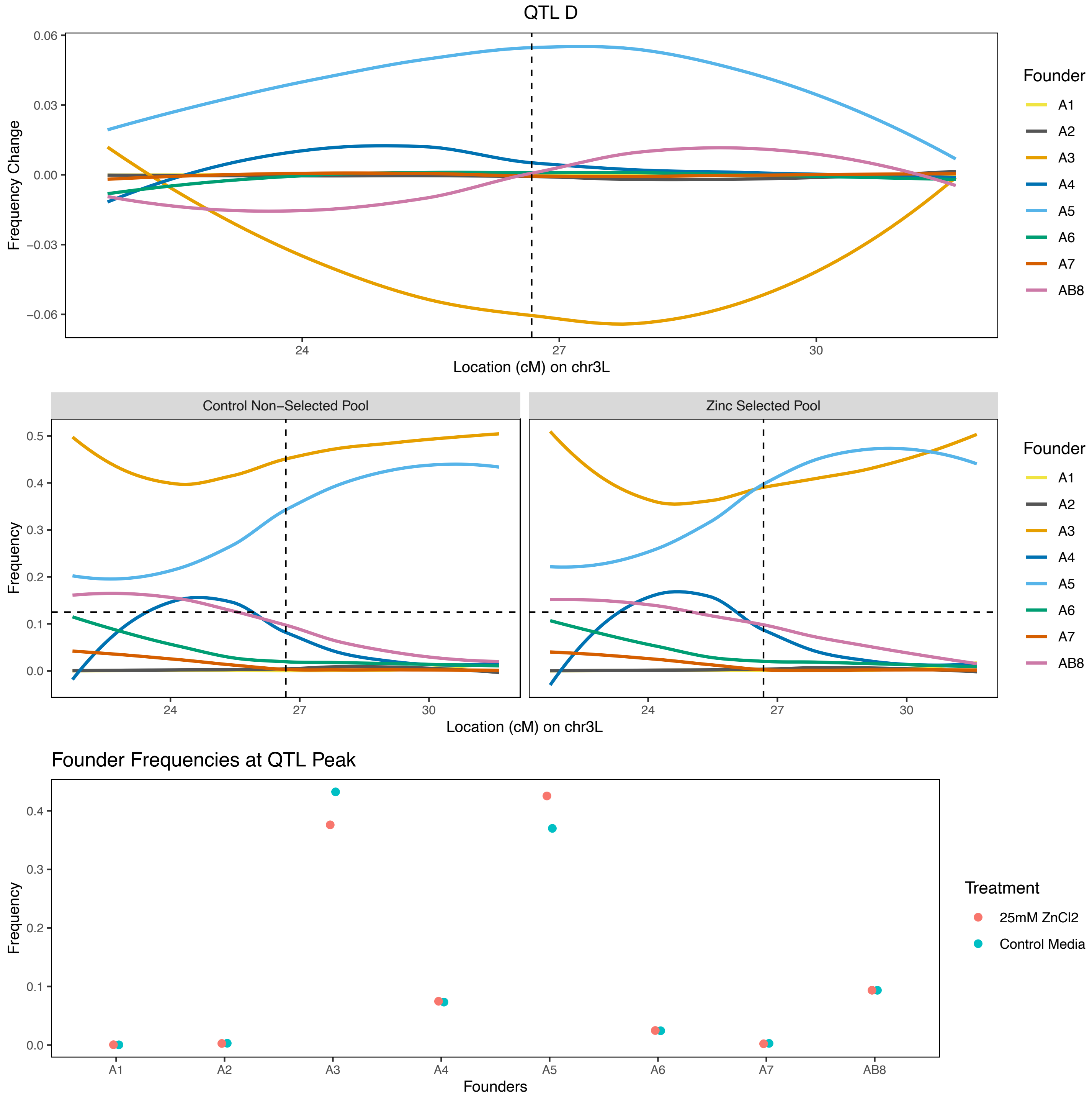

**Supplementary Figure 2:** Contd.

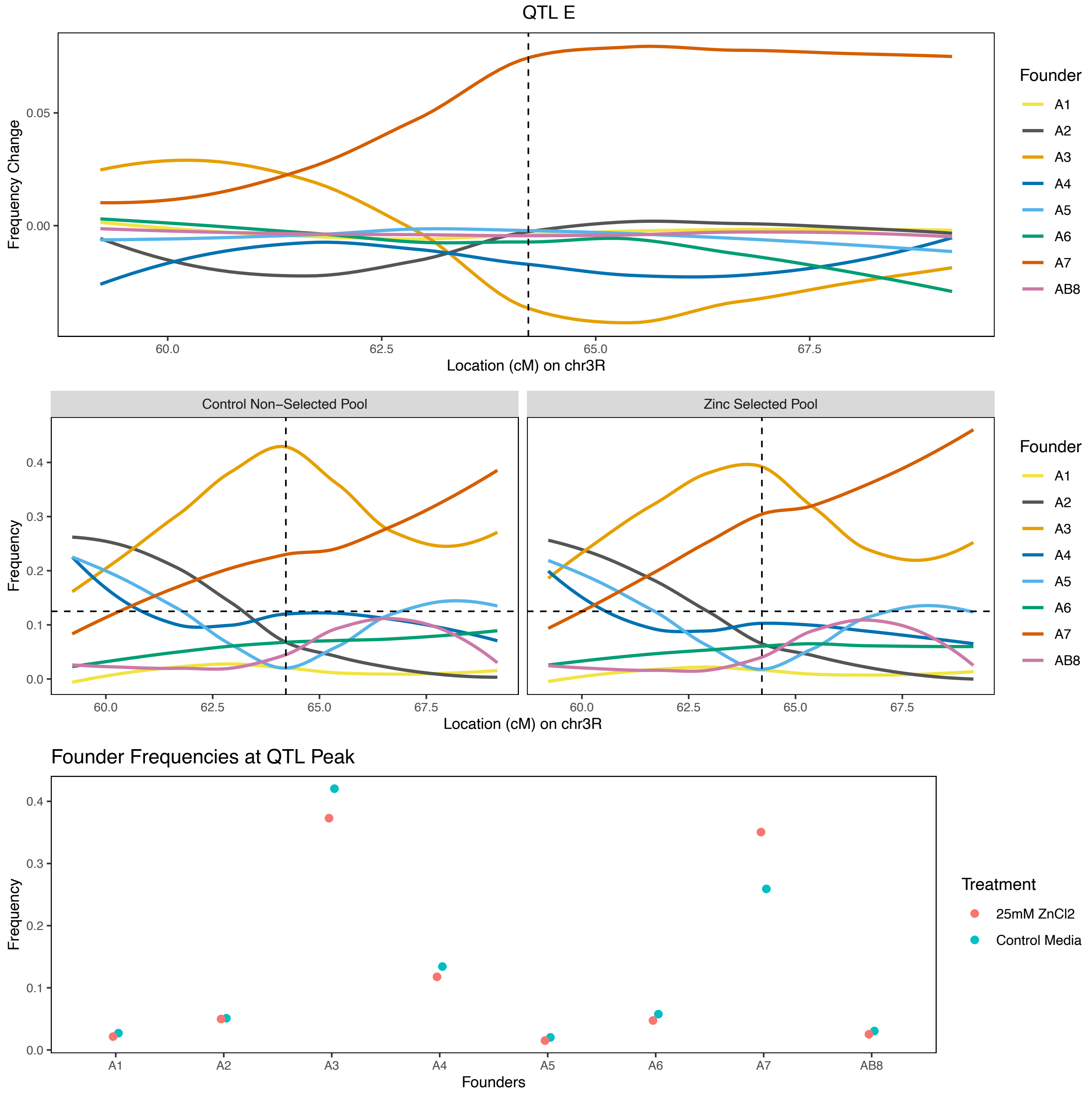

**Supplementary Figure 2:** Contd.

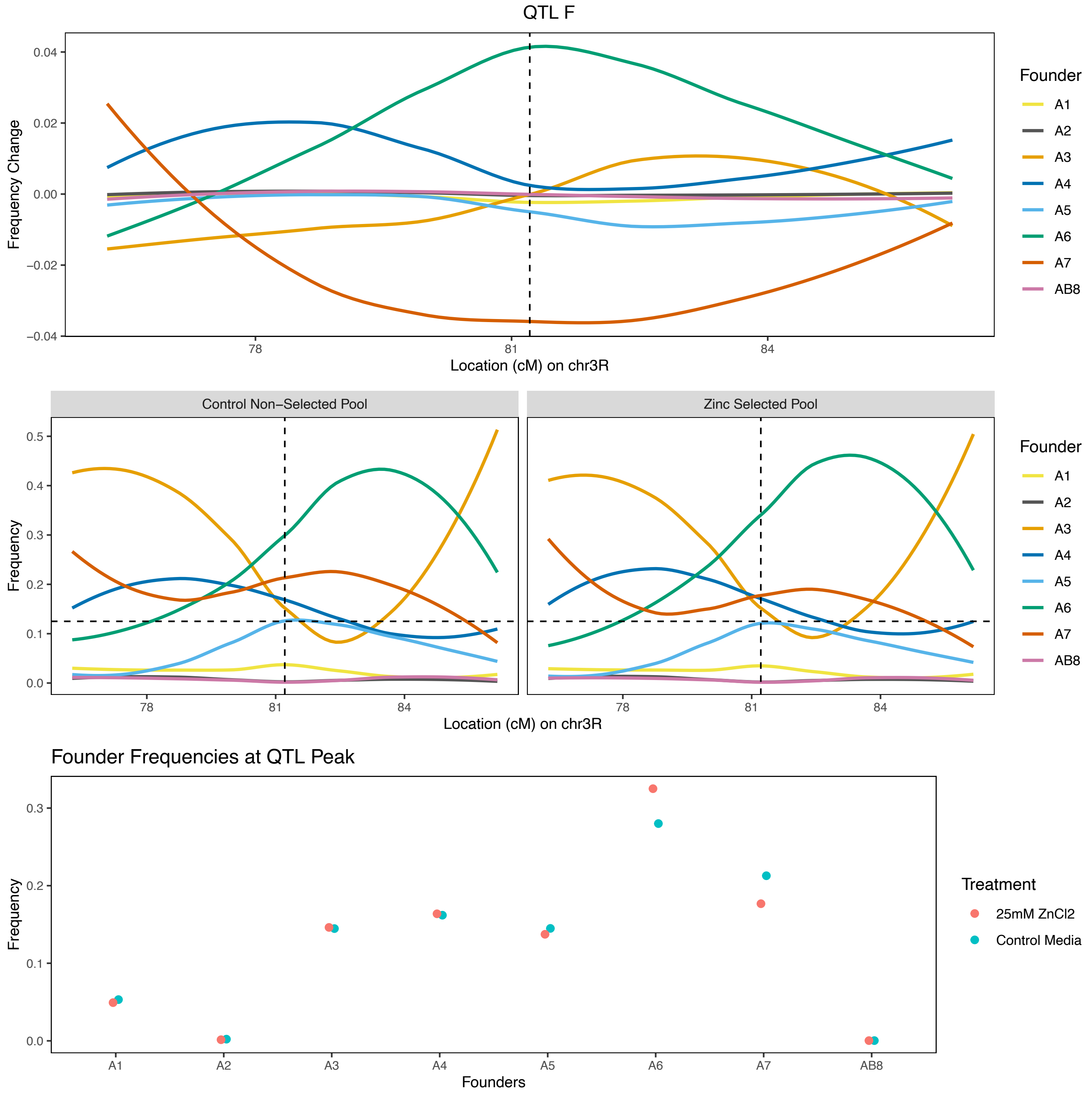

**Supplementary Figure 2:** Contd.

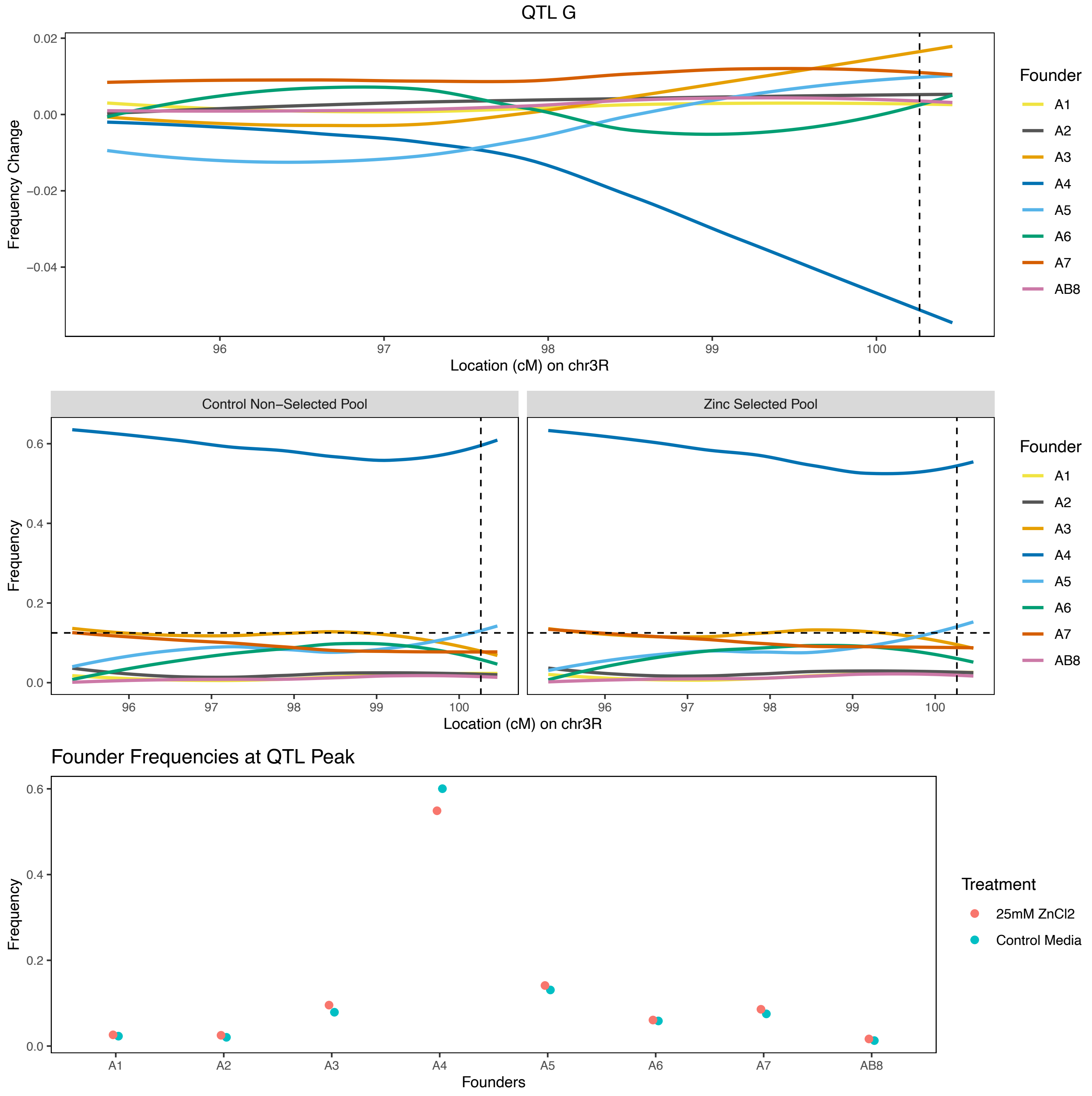

**Supplementary Table 3:** ANOVA tables for the emergence phenotype tested on several sets of animals derived from zinc-selected or control, un-selected populations. We do not consider sex as a factor when examining the emergence phenotype since we do not know the sex of the embryos that were placed in the test vials. Additionally, only the "X-QTL Subpopulation" was assayed over replicate batches, so replicate is not considered in the other two analyses. (Table is associated with Figure 3 in the main text.)

*X-QTL Subpopulation Emergence in Water-control or Zinc-containing Treatments (Figure 3A)*

| Factor | Df | Sum Sq | Mean Sq | F value | Pr(>F) |
|---|---|---|---|---|---|
| Replicate | 1 | 0.00035283 | 0.00035283 | 0.07967825 | 0.77908816 |
| Population | 1 | 0.062398 | 0.062398 | 14.0909992 | 0.00051774 |
| Treatment | 1 | 4.93959025 | 4.93959025 | 1115.48061 | 2.17E-32 |
| Replicate:Population | 1 | 0.00382413 | 0.00382413 | 0.8635825 | 0.35792577 |
| Replicate:Treatment | 1 | 0.00705382 | 0.00705382 | 1.5929253 | 0.21371151 |
| Population:Treatment | 1 | 0.0084848 | 0.0084848 | 1.91607577 | **0.17343227** |
| Replicate:Population:Treatment | 1 | 0.00491813 | 0.00491813 | 1.11063405 | 0.2978301 |
| Residuals | 43 | 0.19041333 | 0.00442822 | nan | nan |

*Semi-inbred Line Emergence in Water-control or Zinc-containing Treatments (Figure 3B)*

| Factor | Df | Sum Sq | Mean Sq | F value | Pr(>F) |
|---|---|---|---|---|---|
| Population | 1 | 0.02962922 | 0.02962922 | 1.06821751 | 0.30279943 |
| Treatment | 1 | 1.95817271 | 1.95817271 | 70.597684 | 1.59E-14 |
| Population:Treatment | 1 | 0.00079139 | 0.00079139 | 0.02853183 | **0.86606302** |
| Residuals | 172 | 4.77077558 | 0.02773707 | nan | nan |

*Phenotyping-Only cohorts Emergence in Water-control or Zinc-containing Treatments (Figure 3C)*

| Factor | Df | Sum Sq | Mean Sq | F value | Pr(>F) |
|---|---|---|---|---|---|
| Population | 1 | 0.04952116 | 0.04952116 | 8.73412322 | 0.00361778 |
| Treatment | 1 | 23.9417856 | 23.9417856 | 4222.64953 | 2.54E-113 |
| Population:Treatment | 1 | 0.03324203 | 0.03324203 | 5.86294704 | **0.016633** |
| Residuals | 153 | 0.86748691 | 0.00566985 | nan | nan |

**Supplementary Table 4:** ANOVA tables for the development time phenotype tested on several sets of animals derived from zinc-selected or control, un-selected populations. Only the "X-QTL Subpopulation" was assayed over replicate batches, so replicate is not considered in the other two analyses. (Associated with Figure 4.)

*X-QTL Subpopulation Development Time in Water-control or Zinc-containing Treatments (Figure 4A)*

| Factor | Df | Sum Sq | Mean Sq | F value | Pr(>F) |
|---|---|---|---|---|---|
| Replicate | 1 | 103.287902 | 103.287902 | 59.9639017 | 3.49E-14 |
| Population | 1 | 3.52536194 | 3.52536194 | 2.04665265 | 0.15299858 |
| Treatment | 1 | 4128.89214 | 4128.89214 | 2397.03275 | 1.42E-225 |
| Sex | 1 | 1.66583598 | 1.66583598 | 0.96710286 | 0.32575201 |
| Replicate:Population | 1 | 18.2856494 | 18.2856494 | 10.6157533 | 0.00117681 |
| Replicate:Treatment | 1 | 2.55154107 | 2.55154107 | 1.4812999 | 0.22399162 |
| Population:Treatment | 1 | 23.269981 | 23.269981 | 13.5094124 | **0.00025599** |
| Replicate:Sex | 1 | 1.32664343 | 1.32664343 | 0.77018426 | 0.38046865 |
| Population:Sex | 1 | 4.85406248 | 4.85406248 | 2.81803117 | 0.09366808 |
| Treatment:Sex | 1 | 0.72456241 | 0.72456241 | 0.42064548 | 0.51683313 |
| Replicate:Population:Treatment | 1 | 4.26393449 | 4.26393449 | 2.47543173 | 0.11610077 |
| Replicate:Population:Sex | 1 | 3.14926669 | 3.14926669 | 1.82831015 | 0.17677482 |
| Replicate:Treatment:Sex | 1 | 5.06302084 | 5.06302084 | 2.93934216 | 0.08689959 |
| Population:Treatment:Sex | 1 | 6.22343735 | 6.22343735 | 3.61302321 | 0.05774953 |
| Replicate:Population:Treatment:Sex | 1 | 13.6957354 | 13.6957354 | 7.95107387 | 0.0049452 |
| Residuals | 683 | 1176.46842 | 1.72250135 | nan | nan |

*Semi-inbred Line Development Time in Water-control or Zinc-containing Treatments (Figure 4B)*

| Factor | Df | Sum Sq | Mean Sq | F value | Pr(>F) |
|---|---|---|---|---|---|
| Population | 1 | 892.791338 | 892.791338 | 245.946935 | 1.81E-53 |
| Treatment | 1 | 20130.7982 | 20130.7982 | 5545.64981 | 0 |
| Sex | 1 | 13.0415602 | 13.0415602 | 3.59270036 | 0.0581234 |
| Population:Treatment | 1 | 411.081608 | 411.081608 | 113.245119 | **5.14E-26** |
| Population:Sex | 1 | 1.78484459 | 1.78484459 | 0.49169054 | 0.48322486 |
| Treatment:Sex | 1 | 1.28305092 | 1.28305092 | 0.35345599 | 0.55220507 |
| Population:Treatment:Sex | 1 | 0.77520082 | 0.77520082 | 0.213553 | 0.64402798 |
| Residuals | 3218 | 11681.3919 | 3.63001611 | nan | nan |

*Phenotyping-Only cohorts Development Time in Water-control or Zinc-containing Treatments (Figure 4C)*

| Factor | Df | Sum Sq | Mean Sq | F value | Pr(>F) |
|---|---|---|---|---|---|
| Population | 1 | 5.24647536 | 5.24647536 | 1.76834605 | 0.18366898 |
| Treatment | 1 | 16004.7828 | 16004.7828 | 5394.47773 | 0 |
| Sex | 1 | 20.7637731 | 20.7637731 | 6.99851493 | 0.00819229 |
| Population:Treatment | 1 | 2.66621997 | 2.66621997 | 0.89866038 | **0.34320316** |
| Population:Sex | 1 | 4.62242874 | 4.62242874 | 1.55800858 | 0.21203613 |
| Treatment:Sex | 1 | 36.2104188 | 36.2104188 | 12.2048702 | 0.00048224 |
| Population:Treatment:Sex | 1 | 0.06832184 | 0.06832184 | 0.02302816 | 0.87939216 |
| Residuals | 3695 | 10962.6317 | 2.96688273 | nan | nan |

**Supplementary Figure 3:** Variation in phenotype among RNAi control genotypes.

As controls for RNAi knockdowns we examined emergence and development time for 4 different controls; UAS-GFP, UAS-Luciferase, and strains that carry empty attP2 or attP40 docking sites. Parental inbred control strains were crossed to the *mex1*-GAL4 strain, and the F1 genotypes (e.g., *mex1*-GAL4:UAS-GFP) were tested on either water control or zinc-containing media. Notably, *mex1*-GAL4:attP2 (highlighted in blue in the figure) exhibited substantially lower emergence in the $ZnCl_2$ treatment (left panel), and was the slowest developing of all 4 control genotypes on both media types (right panel). See also Supplementary Table 5. Note the inverted *y*-axis used for the development time (right) panel. This was done such that a visibly negative slope for both emergence *and* development time yielded the same inference; zinc treatment led to a worse outcome (reduced emergence or longer development time.) Points are phenotype means for a given genotype/treatment, and vertical bars are 95% confidence intervals.

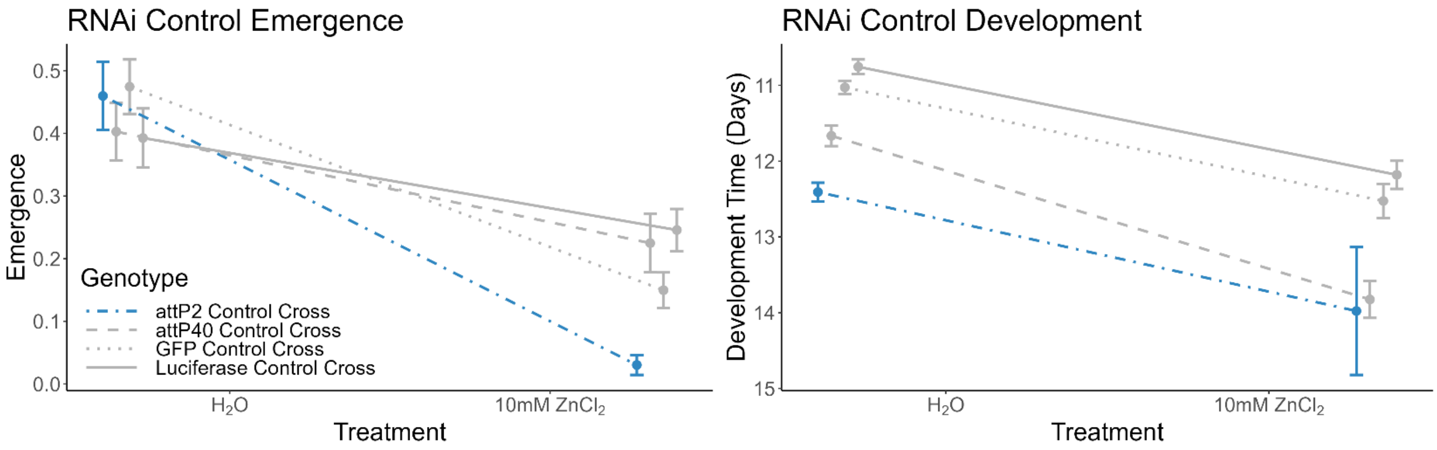

**Supplementary Table 5:** Summary of variation in phenotype among RNAi control genotypes.

| Control Genotype * | Water Control Treatment | | | | 10mM $ZnCl_2$ | | | |
|---|---|---|---|---|---|---|---|---|
| | Total Vials ** | Total Flies Emerged | Average Emergence | Average Development Time | Total Vials ** | Total Flies Emerged | Average Emergence | Average Development Time |
| attP2 | 29 | 665 | 0.460 | **12.4** | 29 | 44 | **0.030** | **14.0** |
| attP40 | 29 | 580 | 0.402 | 11.7 | 29 | 326 | 0.225 | 13.8 |
| UAS-GFP | 29 | 686 | 0.474 | 11.0 | 29 | 217 | 0.150 | 12.5 |
| UAS-Luc | 29 | 567 | 0.393 | 10.8 | 28 | 344 | 0.246 | 12.2 |

* Genotypes tested are F1 heterozygotes between the *mex1*-GAL4 driver strain and the listed UAS/control strain.

** Each vial initially contains 50 eggs.

**Supplementary Table 6:** Full ANOVA tables for each comparison between an RNAi knockdown genotype and each genetic control for both emergence and development time. Of interest for each test is the significance of the genotype-by-treatment interaction, since this reflects any difference between knockdown and controls in their response to zinc treatment. In the table the "Phenotype" column indicates whether the test is for emergence or development time, "Control" indicates which genetic control the RNAi knockdown is compared to, "Gene_KD_UAS_Line" gives the gene ID and the Bloomington *Drosophila* Stock Center (BDSC) ID for the UAS-RNAi strain used, and "UAS_Chromosome" defines which chromosome (chromosome 2 or 3) the UAS-RNAi transgene resides on. Note that due to very low numbers of animals for certain genotype/treatment/sex combinations some coefficients could not be estimated, and not all possible interactions were tested. (Associated with Supplementary Figure 4 and Table 2.)

*Due to its length, Supplementary Table 6 is available for separate download as a *.csv file.*

**Supplementary Figure 4:** Effects of RNAi expression knockdown on emergence and development time.

Each of the 18 figures on the next few pages focuses on a single gene, and has 2 panels; the right panel shows the fraction of egg-to-adult emergence, the left panel highlights development time. Note the inverted *y*-axis used for the development time (right) panel. This was done such that a visibly negative slope for both emergence *and* development time yields the same inference; zinc treatment leads to a worse outcome (reduced emergence or longer development time.) Each of the plots shows the exact same data for the 4 control genotypes (*mex1*-GAL4:attP2, *mex1*-GAL4:attP40, *mex1*-GAL4:UAS-GFP, and *mex1*-GAL4:UAS-Luc) in various gray lines, along with the data for *mex1*-GAL4:UAS-RNAi genotype targeting a specific gene (in red or blue; when we targeted the same gene with two different RNAi transgenes, both are shown on the same plot). Points are means across all vials measured per genotype per treatment, while vertical bars are 95% confidence intervals. Our analyses are focused on identifying significant genotype-by-treatment effects of RNAi knockdown (i.e., is the effect of treatment different for the RNAi knockdown and genetic control genotypes), which is seen in these plots via a different slope. See in-text Table 2 for a summary of the statistical analyses, and Supplementary Table 6 for full details of the ANOVAs.

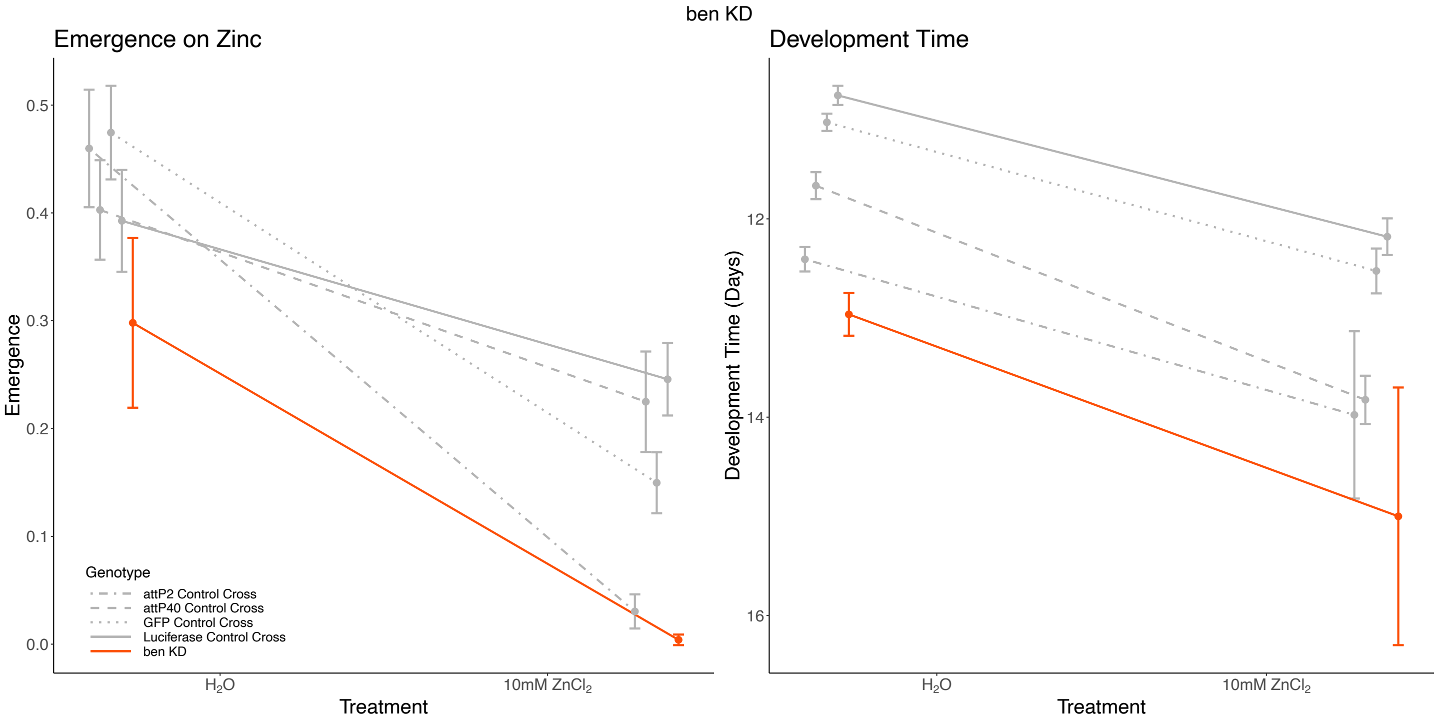

**Supplementary Figure 4:** Contd.

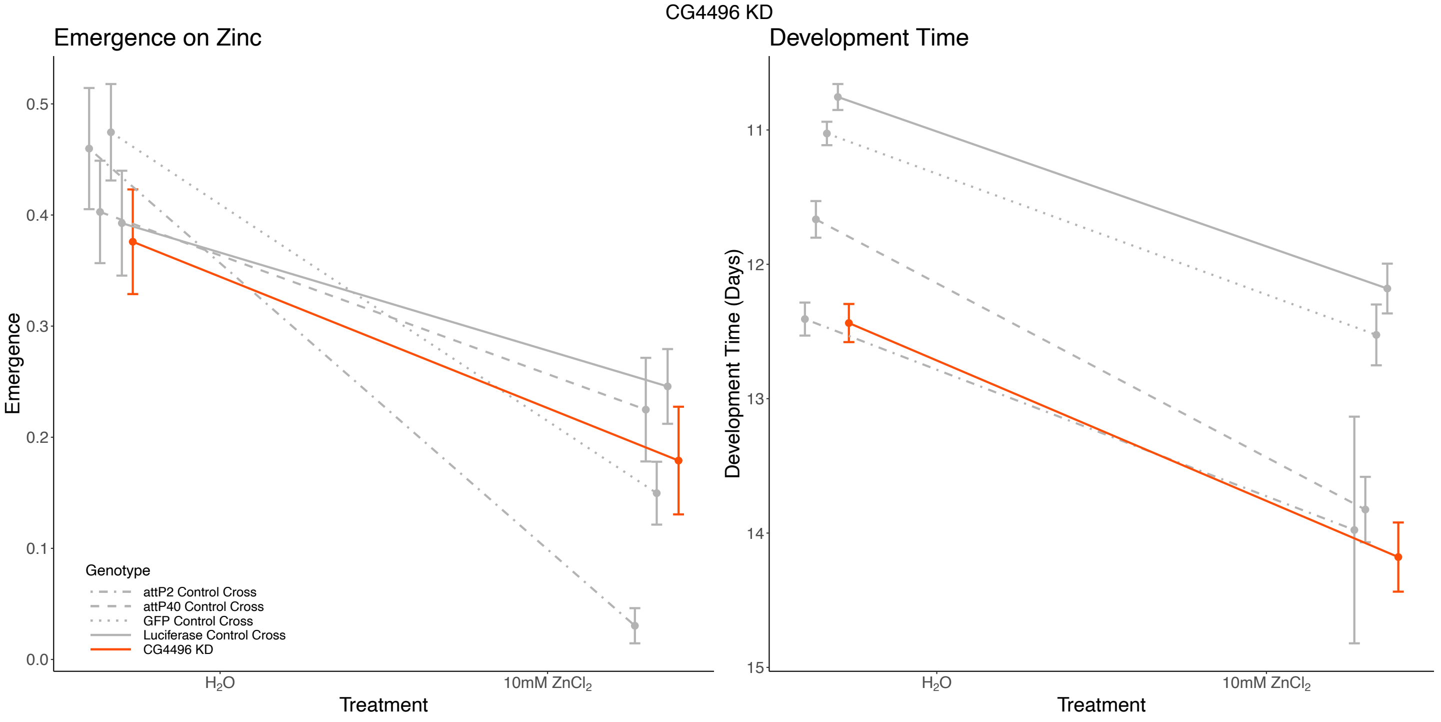


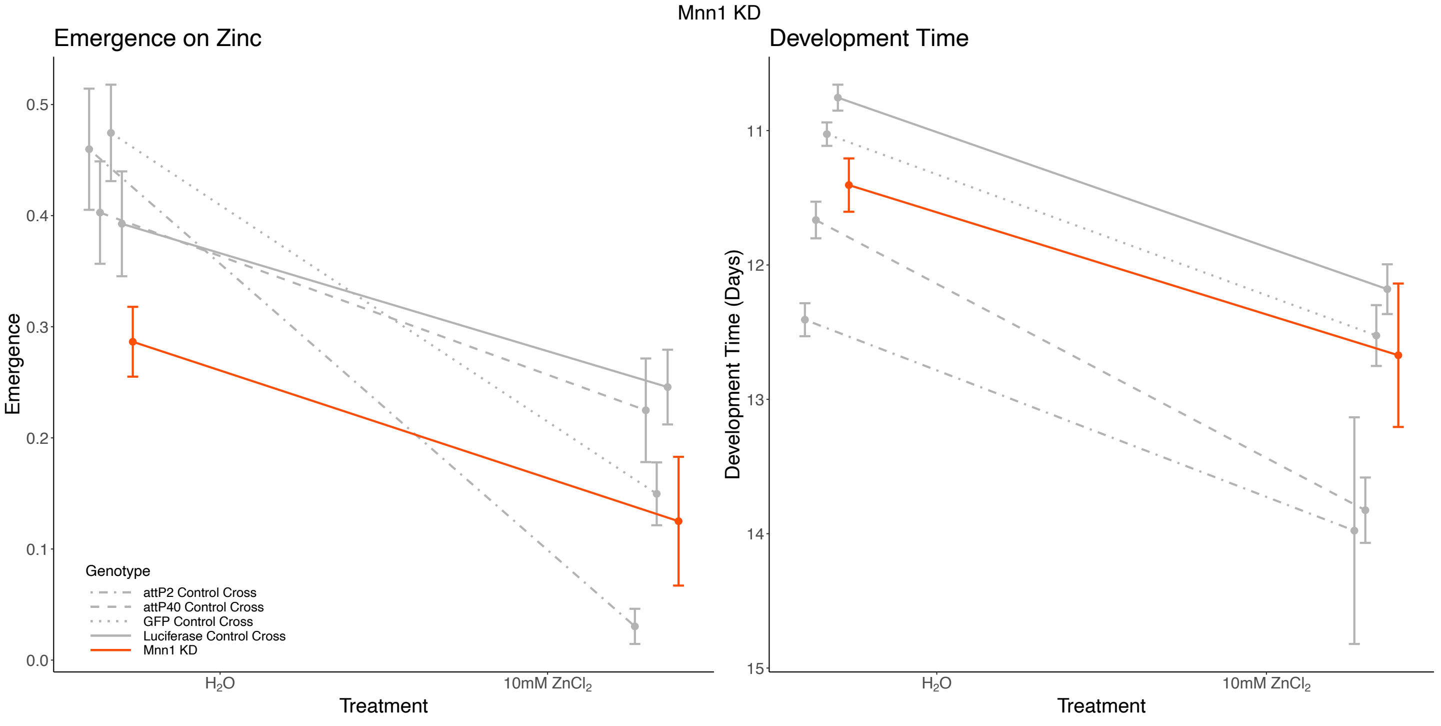

**Supplementary Figure 4:** Contd.

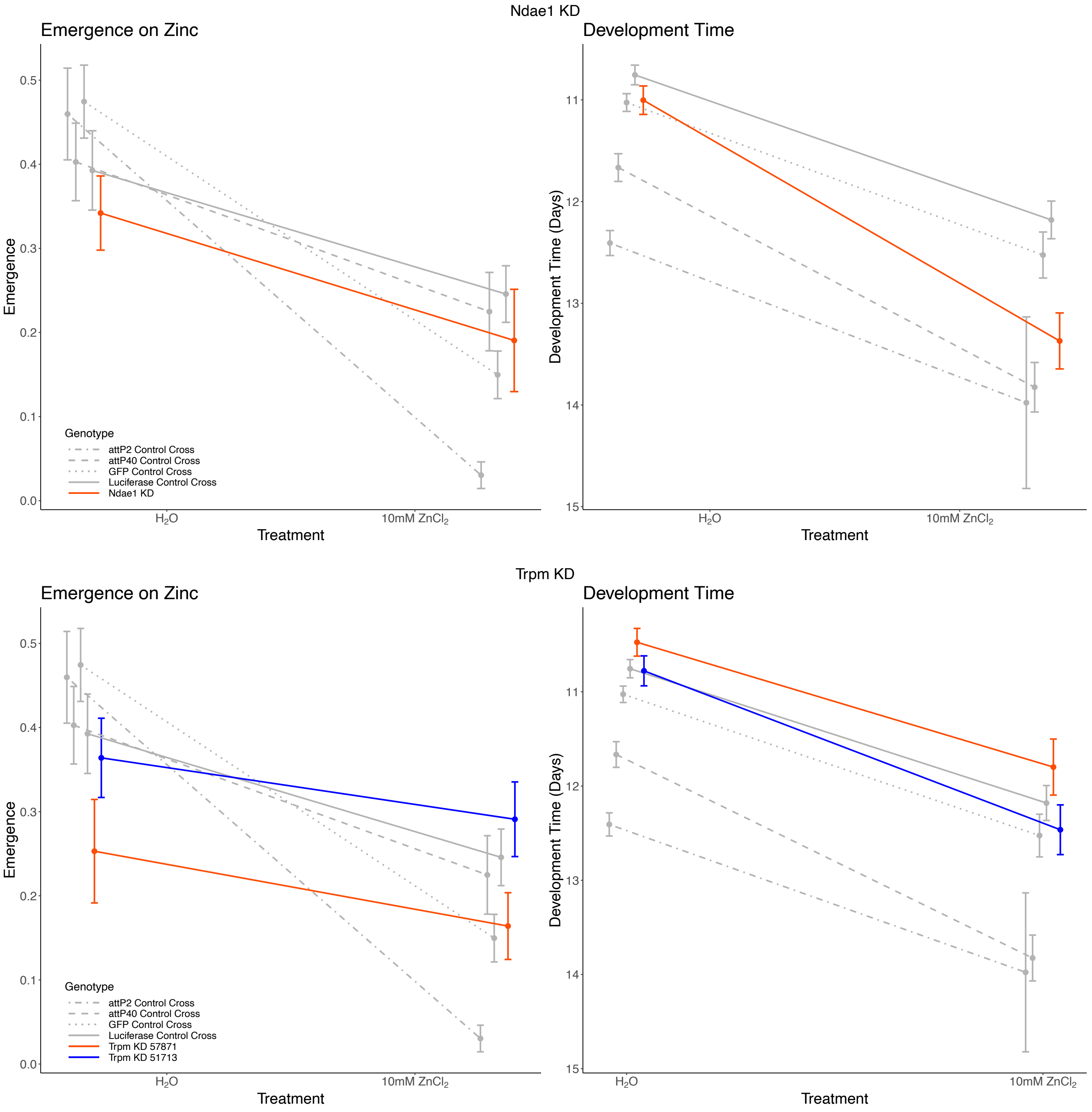

**Supplementary Figure 4:** Contd.

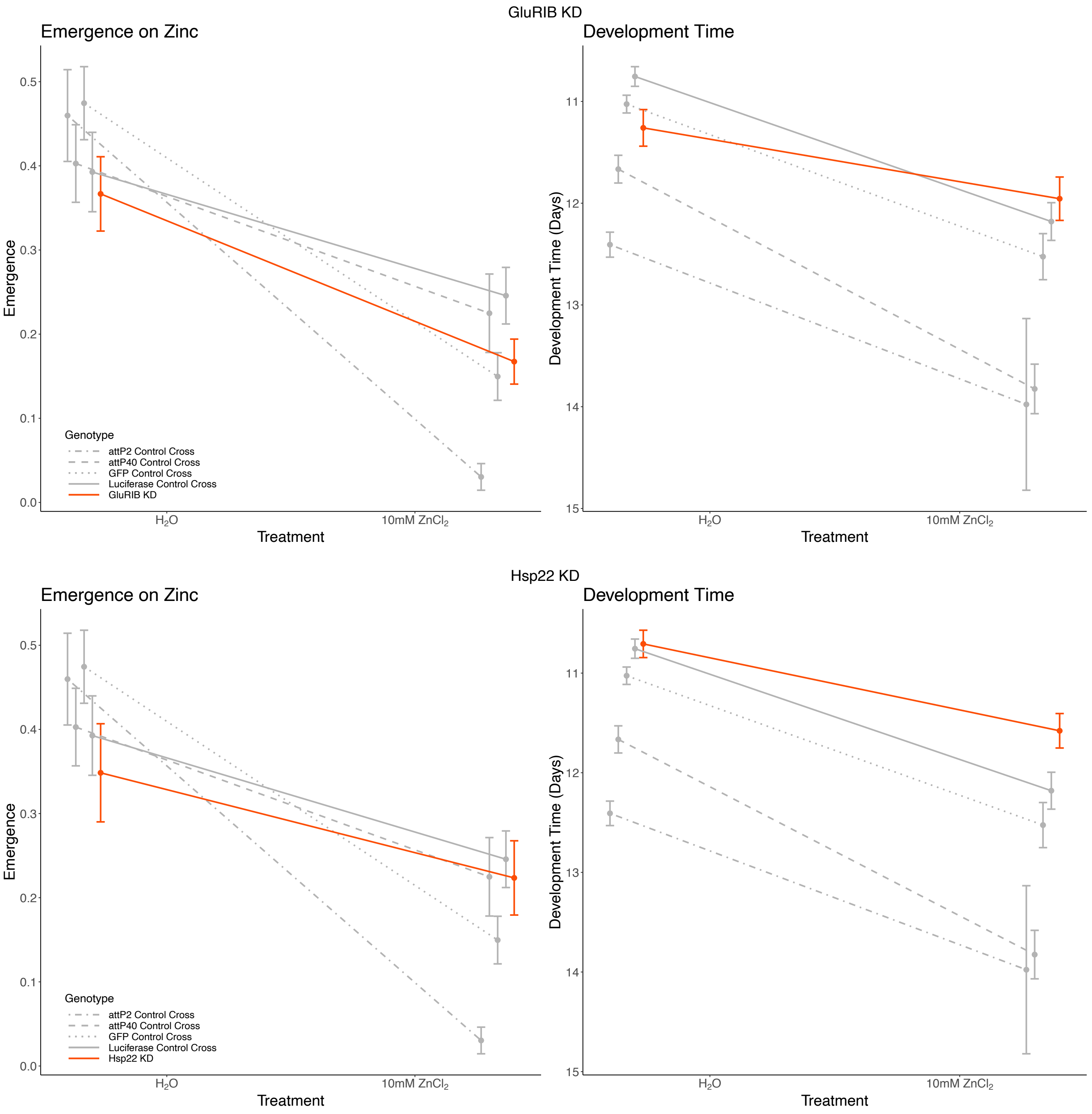

**Supplementary Figure 4:** Contd.

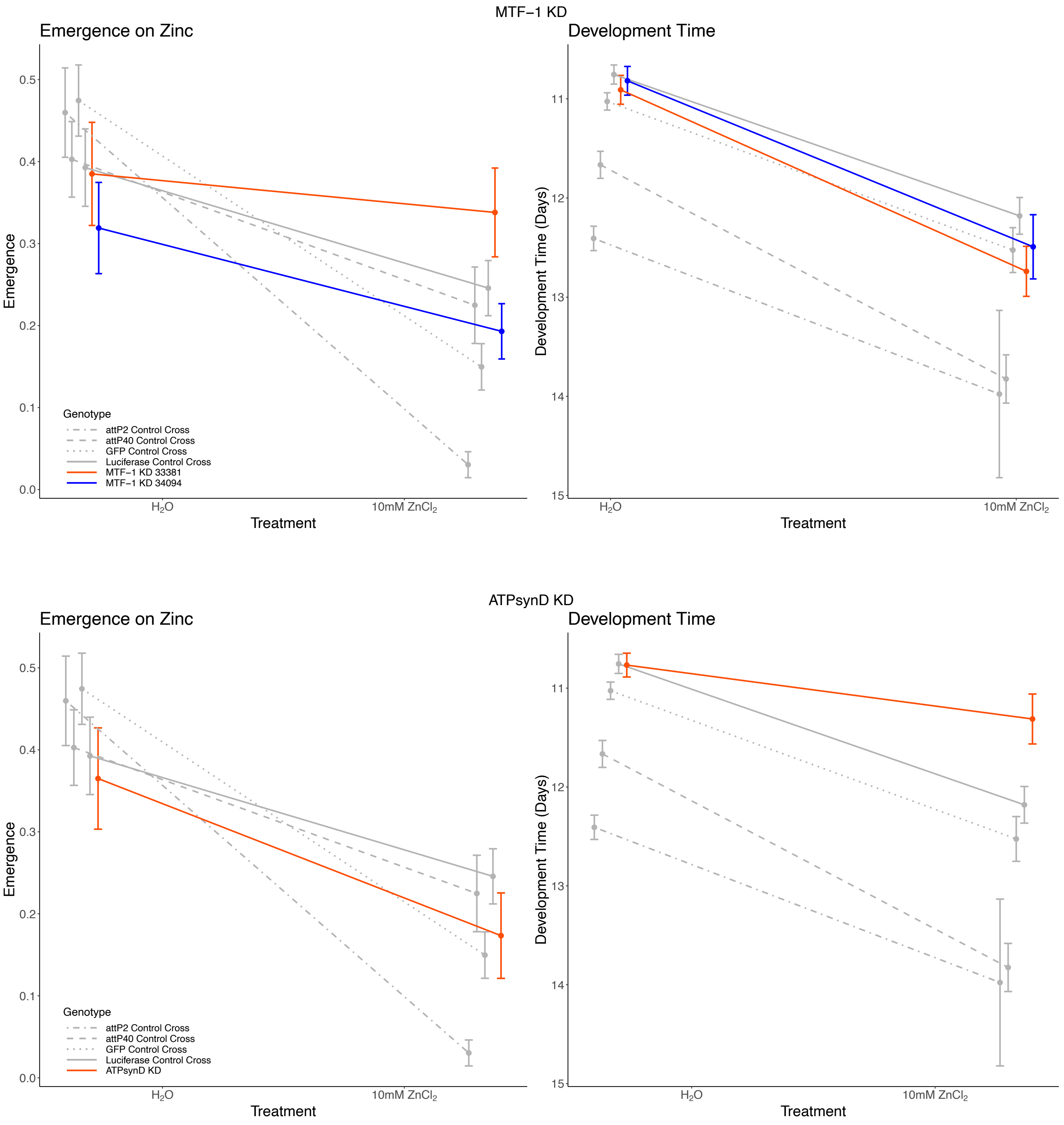

**Supplementary Figure 4:** Contd.

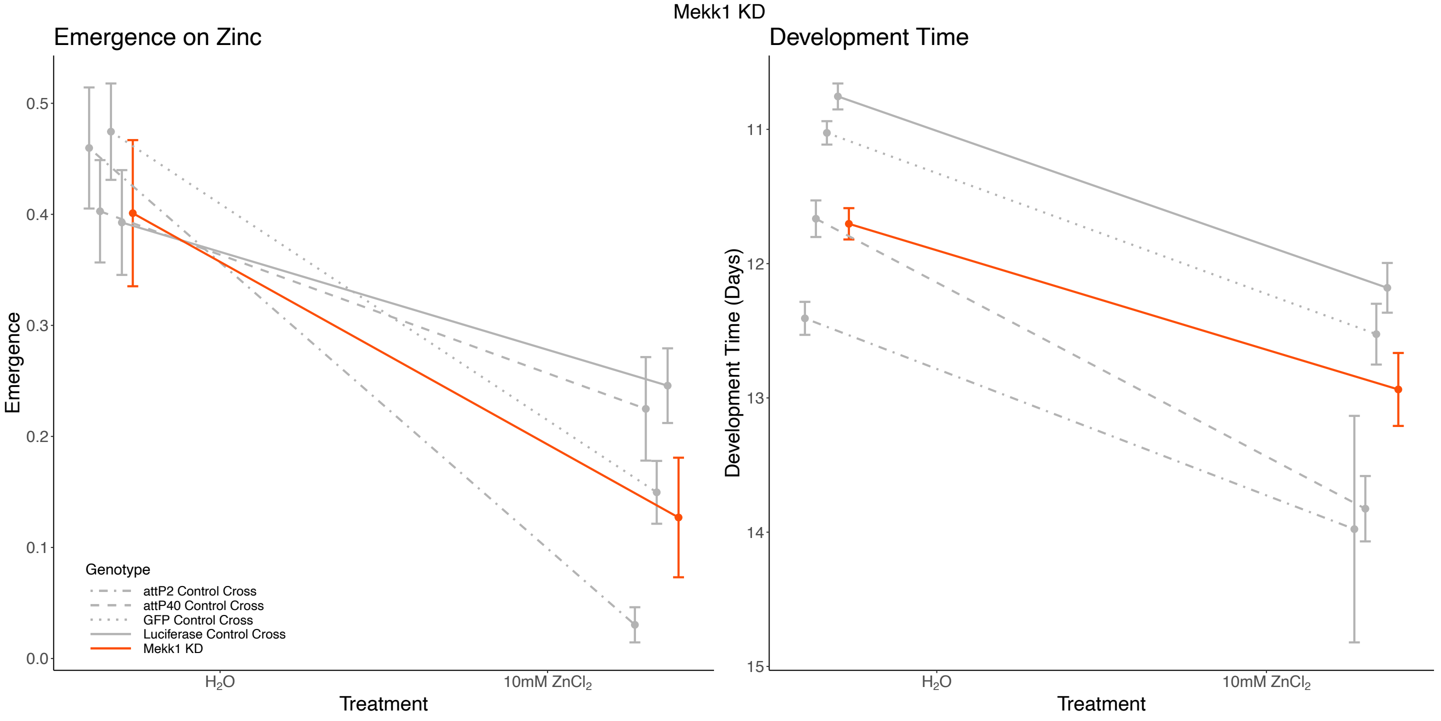


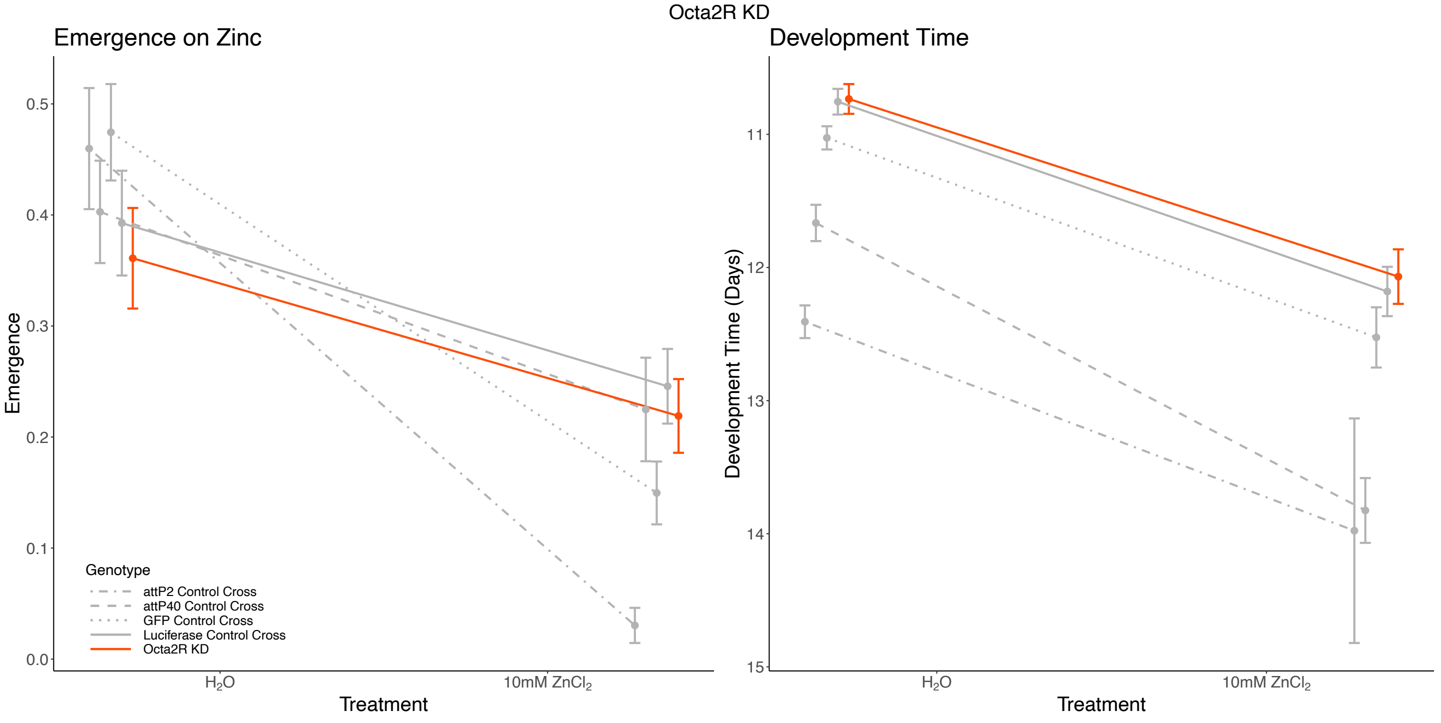

**Supplementary Figure 4:** Contd.

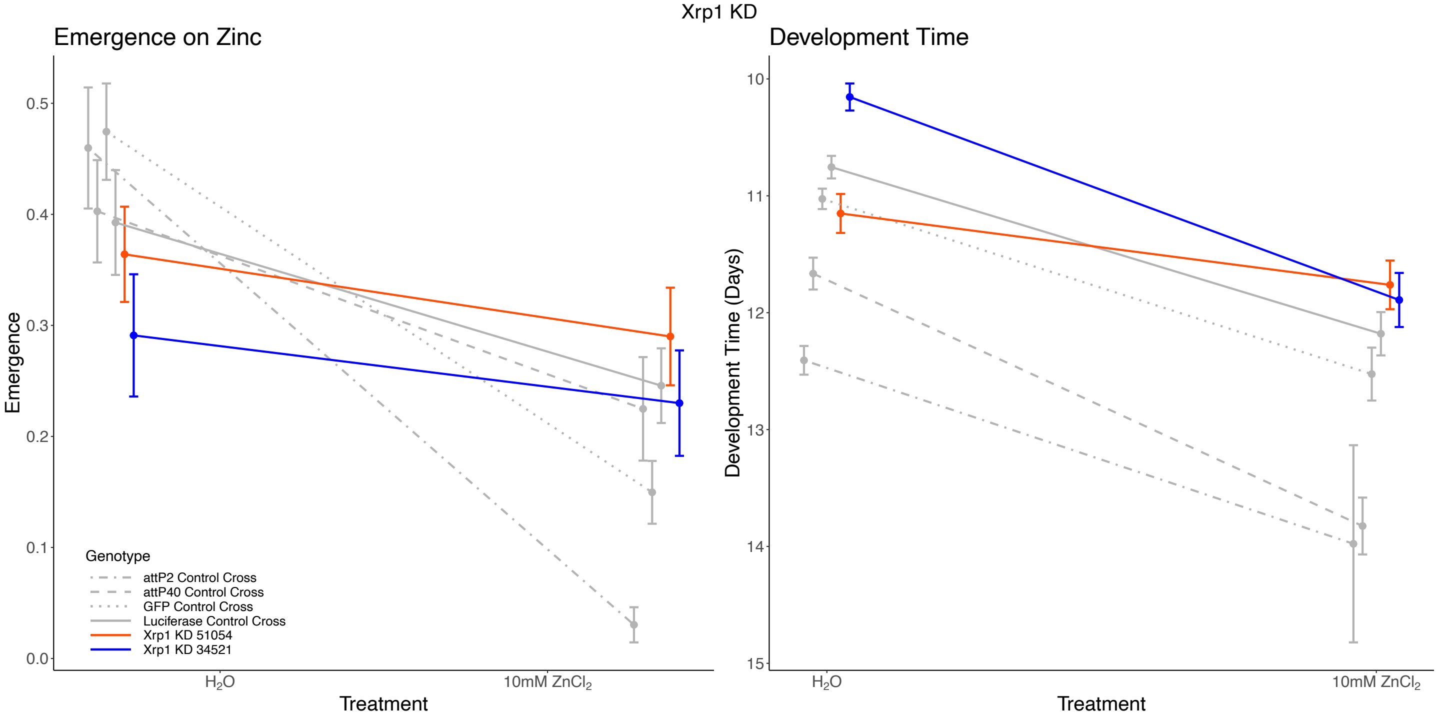


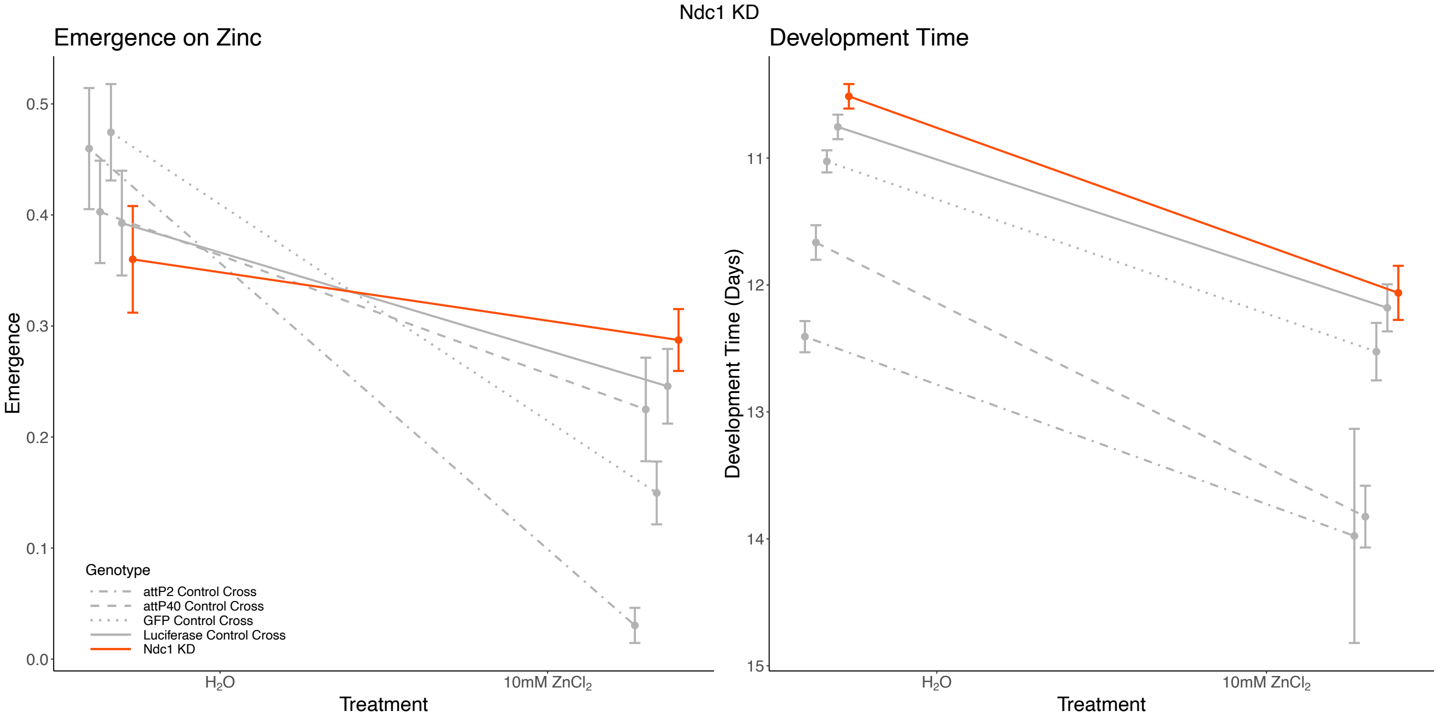

**Supplementary Figure 4:** Contd.

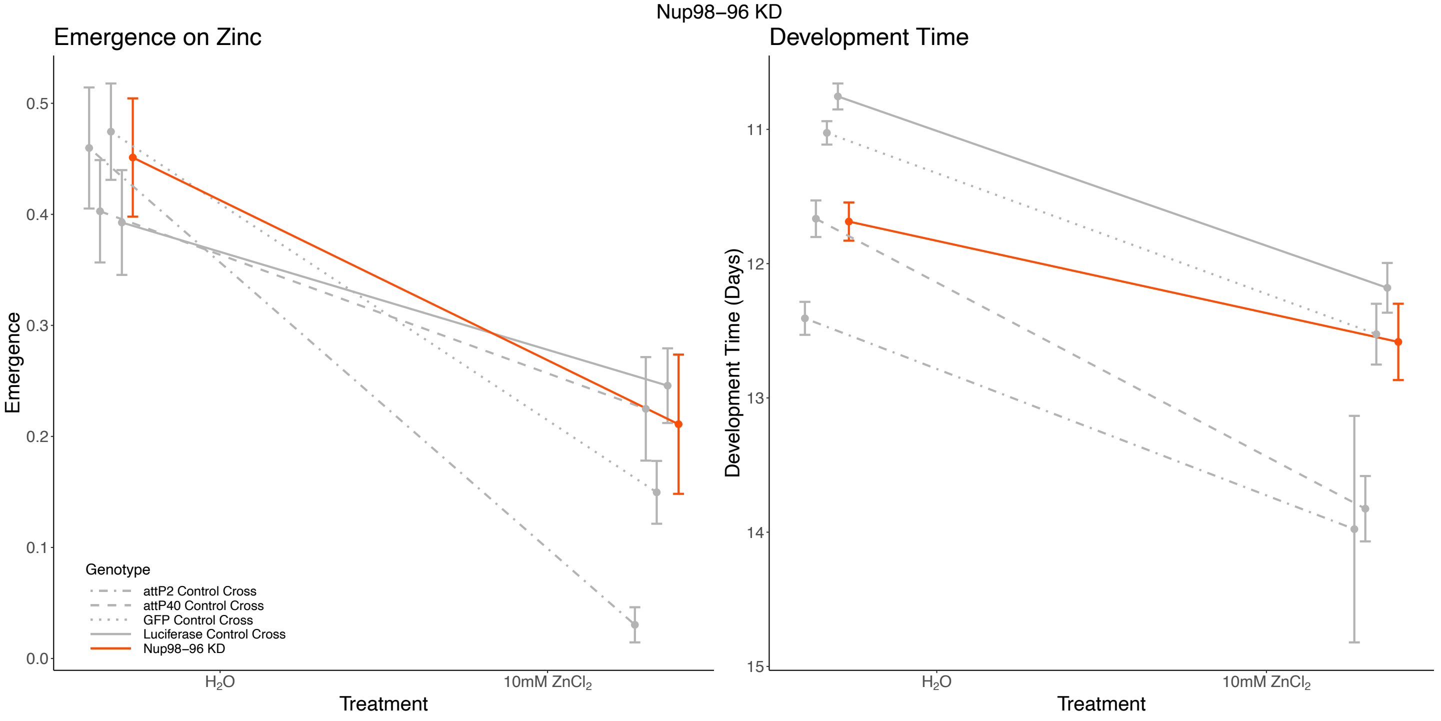


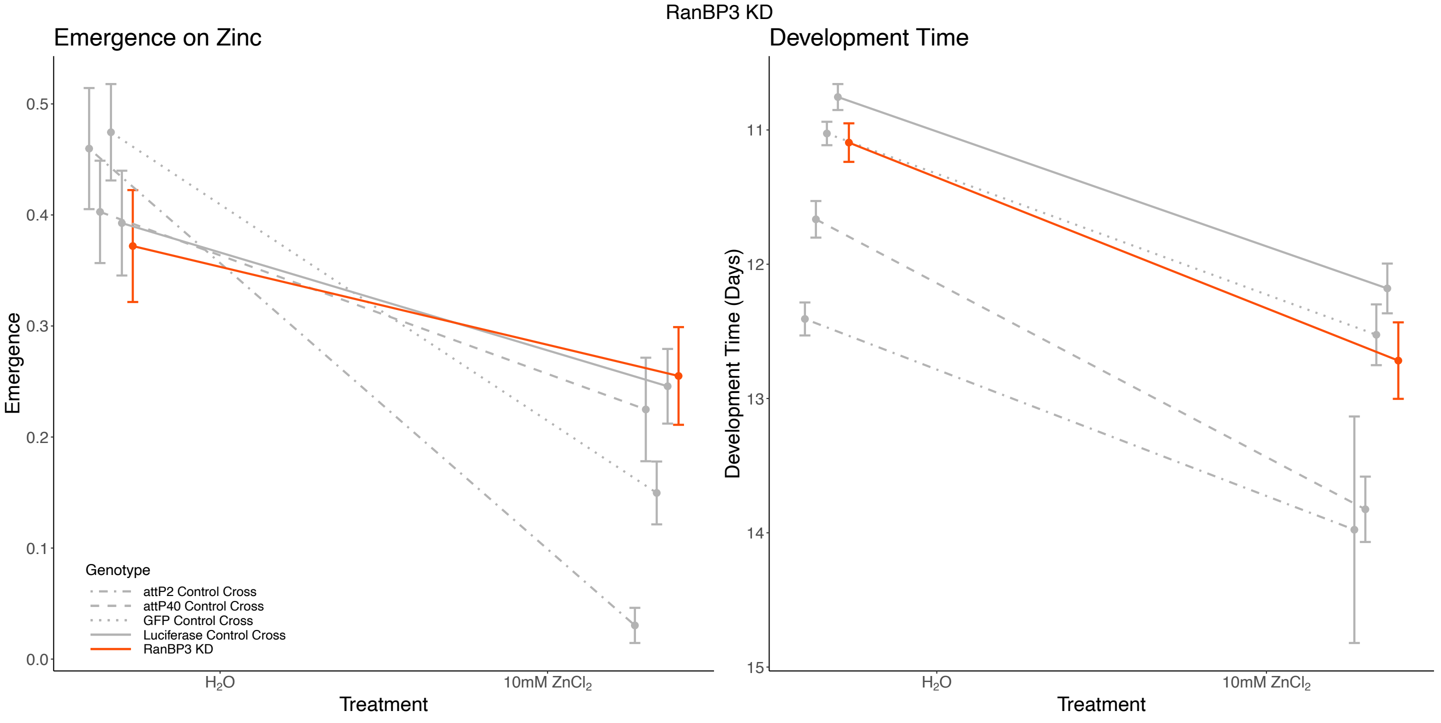

**Supplementary Figure 4:** Contd.

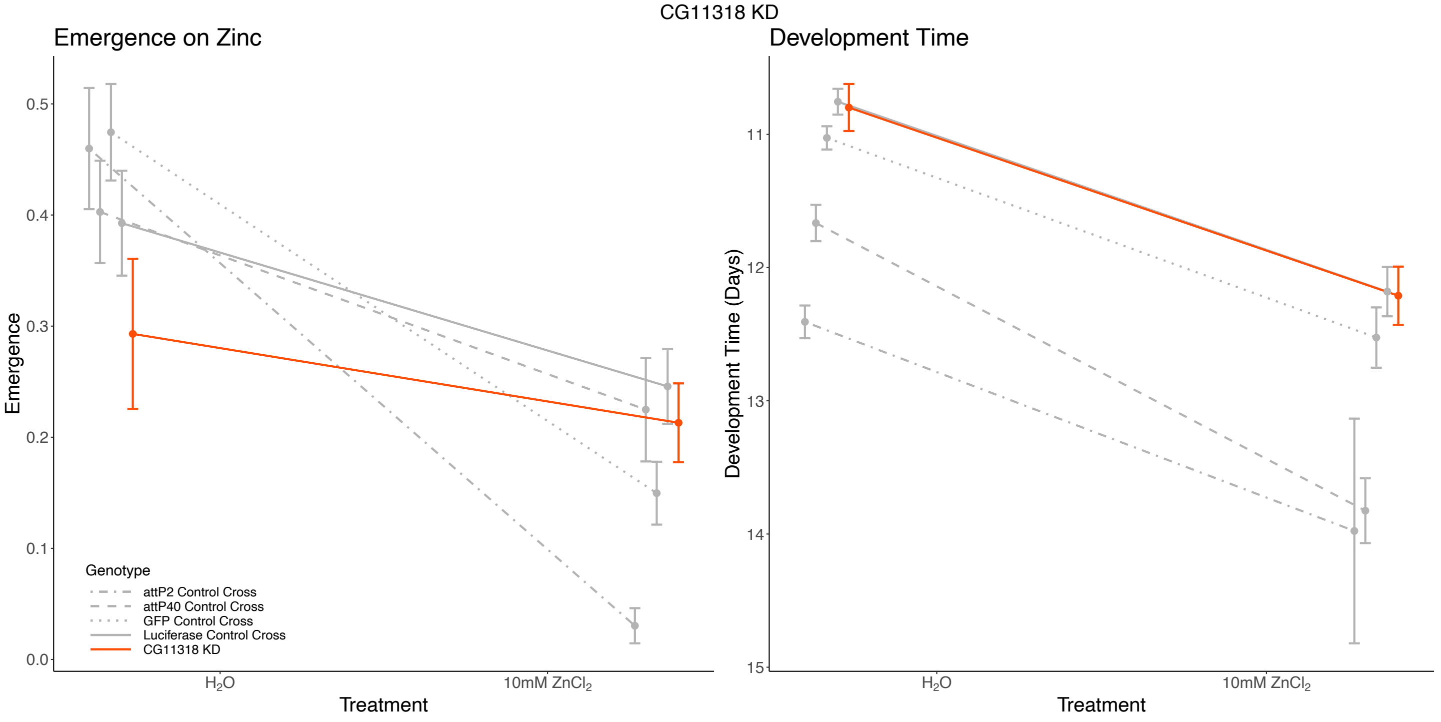


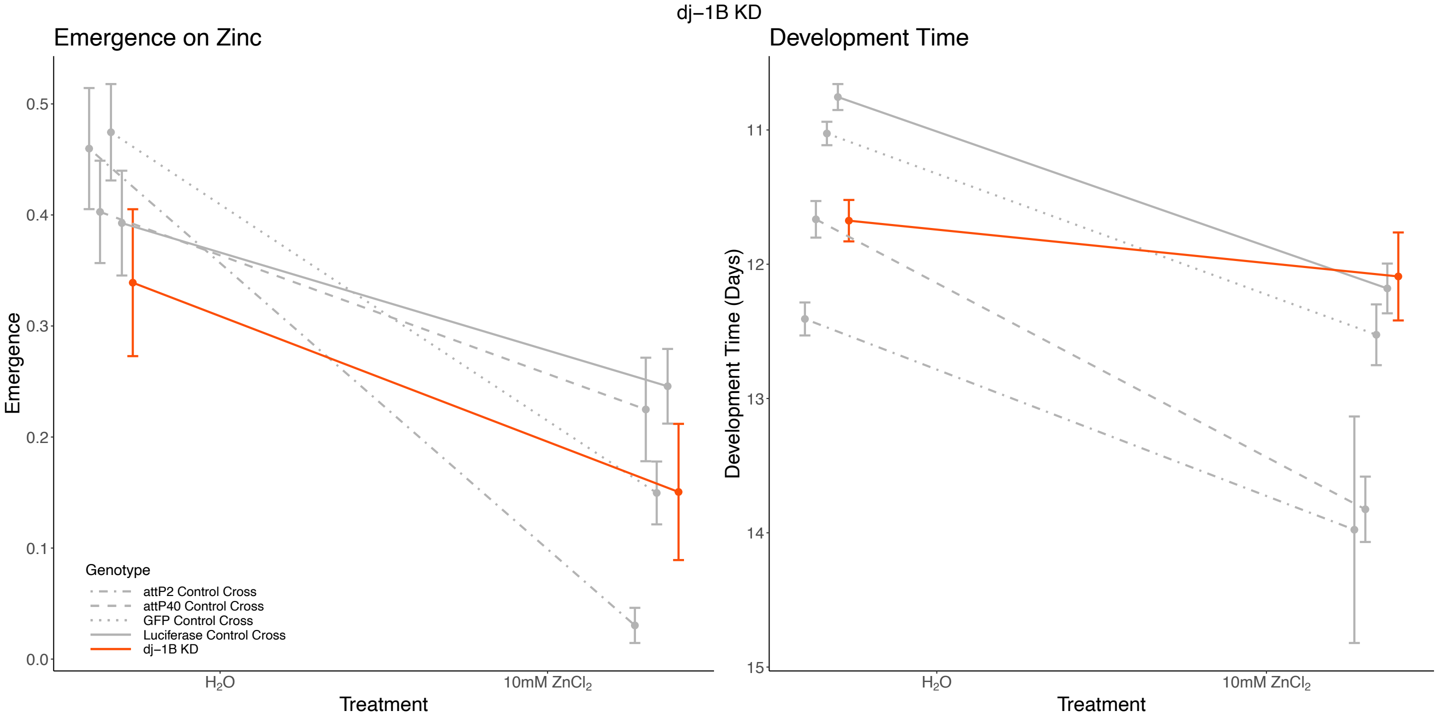

**Supplementary Figure 4:** Contd.

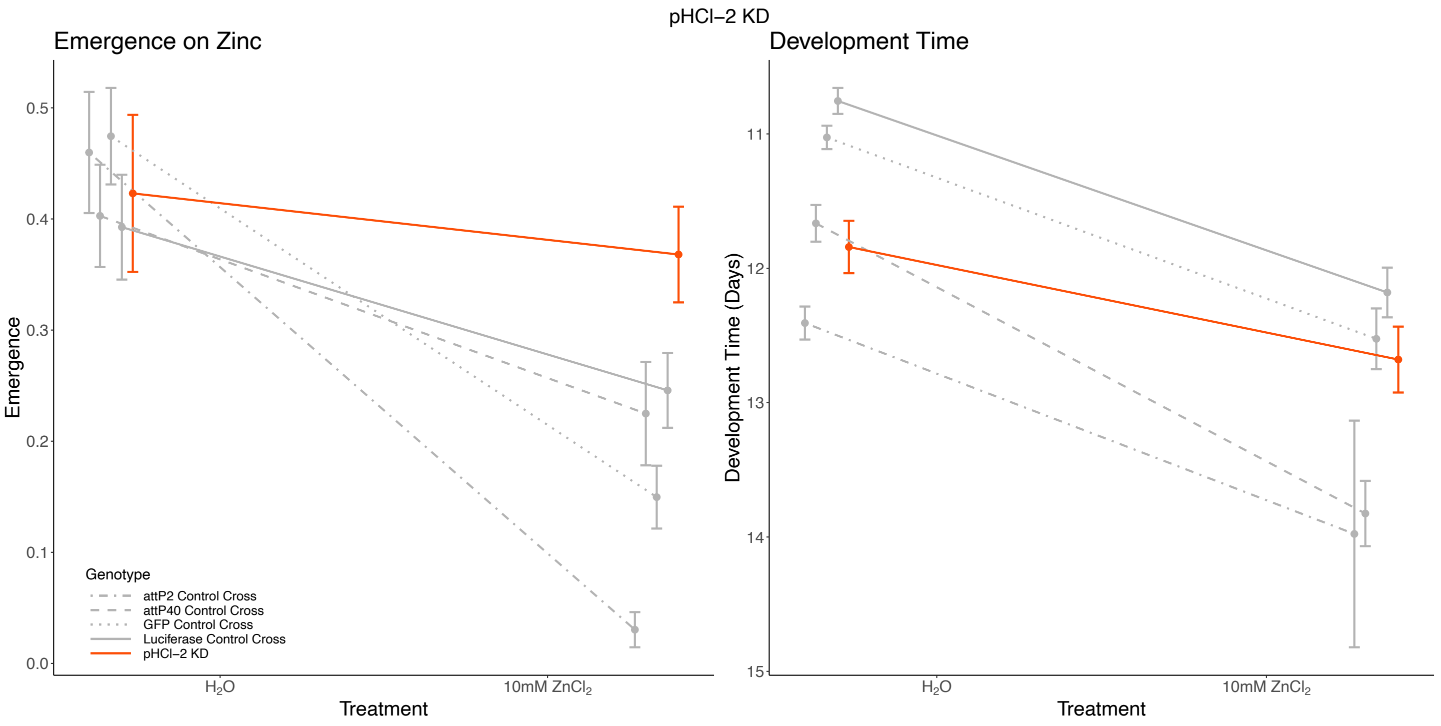

**Supplementary Figure 5:** Adult zinc toxicity.

Adult females from the X-QTL subpopulations (R9C, R9Z, R10C, R10Z) and from the phenotyping-only cohort were raised in normal lab media, and placed into vials with instant media reconstituted with 100mM $ZnCl_2$. The number of dead flies in a vial was recorded each day untill all flies had died. Below we plot the mean time of death and 95% confidence interval for each sample of flies. Adults from the zinc-selected populations either showed no difference in adult resistance, or were slightly less resistant to zinc than non-selected control populations. (We note that flies from each test population were also put in vials reconstituted with water rather than a zinc chloride solution, and no flies died in these vials over the course of the assay).

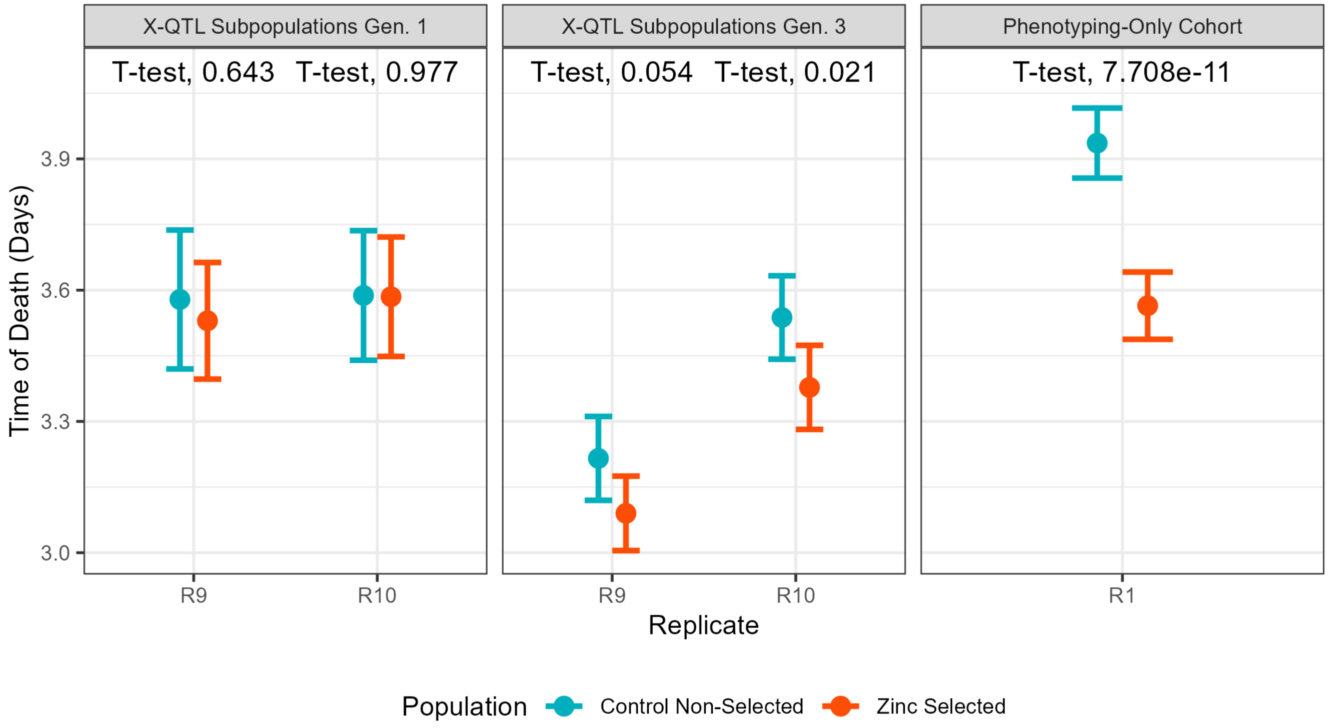

**Supplementary Figure 6:** Emergence rates of flies from media containing different metal salts.

For the phenotyping-only cohorts and the semi-inbred lines, we collected multiple replicate vials of 50 eggs. Vials contained instant media reconstituted either with water or with a different metal solution (25 or 10mM $ZnCl_2$, 0.2mM $CdCl_2$, or 2mM $CuSO_4$). We counted the number of adults emerging from each vial, and in the panels below points represent vial-specific emergence fractions. (A) Phenotyping-only cohorts. We generated outbred animals in an analogous fashion to our X-QTL design. The zinc-selected cohorts showed higher emergence on zinc media than did non-selected controls ($p = 0.017$), but lower emergence on cadmium media ($p < 1 \times 10^{-5}$), and no difference on copper ($p > 0.05$). (B) Semi-inbred Lines. Strains derived from the R9 and R10 X-QTL control and zinc-selected cohorts showed no significant population-by-treatment interaction for any metal. (The $H_2O$ and $ZnCl_2$ data is identical to that presented in Figure 3B and 3C).

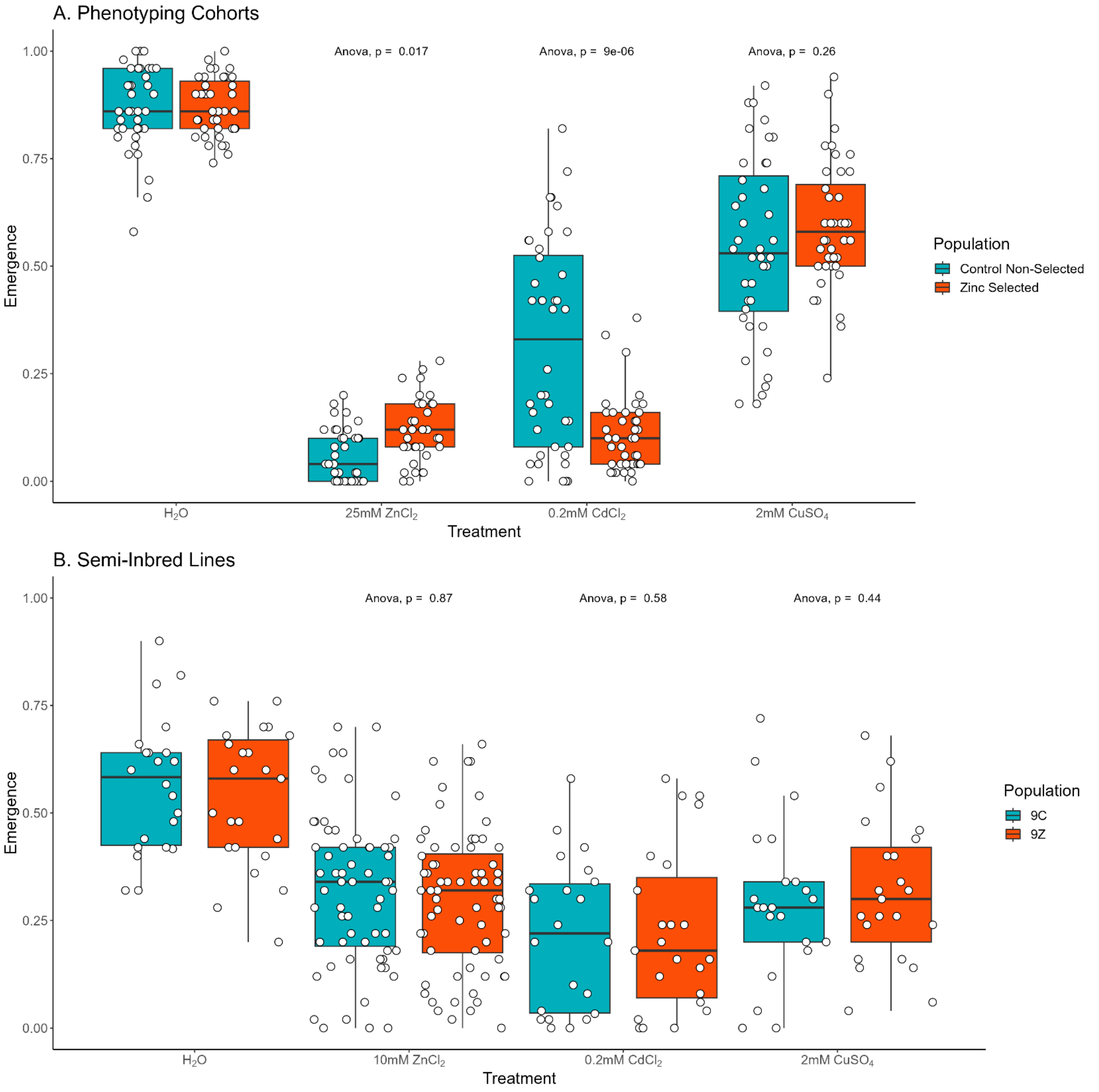

**Supplementary Figure 7:** Development time of flies in media containing different metal salts.

For the phenotyping-only cohorts and the semi-inbred lines, we collected multiple replicate vials of 50 eggs. Vials contained instant media reconstituted either with water or with a different metal solution (25 or 10mM $ZnCl_2$, 0.2mM $CdCl_2$, or 2mM $CuSO_4$). We measured development time from egg collection to adult emergence, and the points in the figures below represent development time of individual animals. (A) Phenotyping-only cohorts. We generated outbred animals in an analogous fashion to our X-QTL design. We see no difference in development between zinc-selected and control animals on zinc media ($p = 0.34$), while zinc-selected animals experience greater developmental delays due to both cadmium ($p < 1 \times 10^{-59}$) and copper ($p < 1 \times 10^{-11}$). (B) Semi-inbred Lines. Strains derived from the R9 and R10 X-QTL control and zinc-selected cohorts showed a significant population-by-treatment interaction effect for zinc, but not for cadmium or copper. (The $H_2O$ and $ZnCl_2$ data is identical to that presented in Figure 4B and 4C).

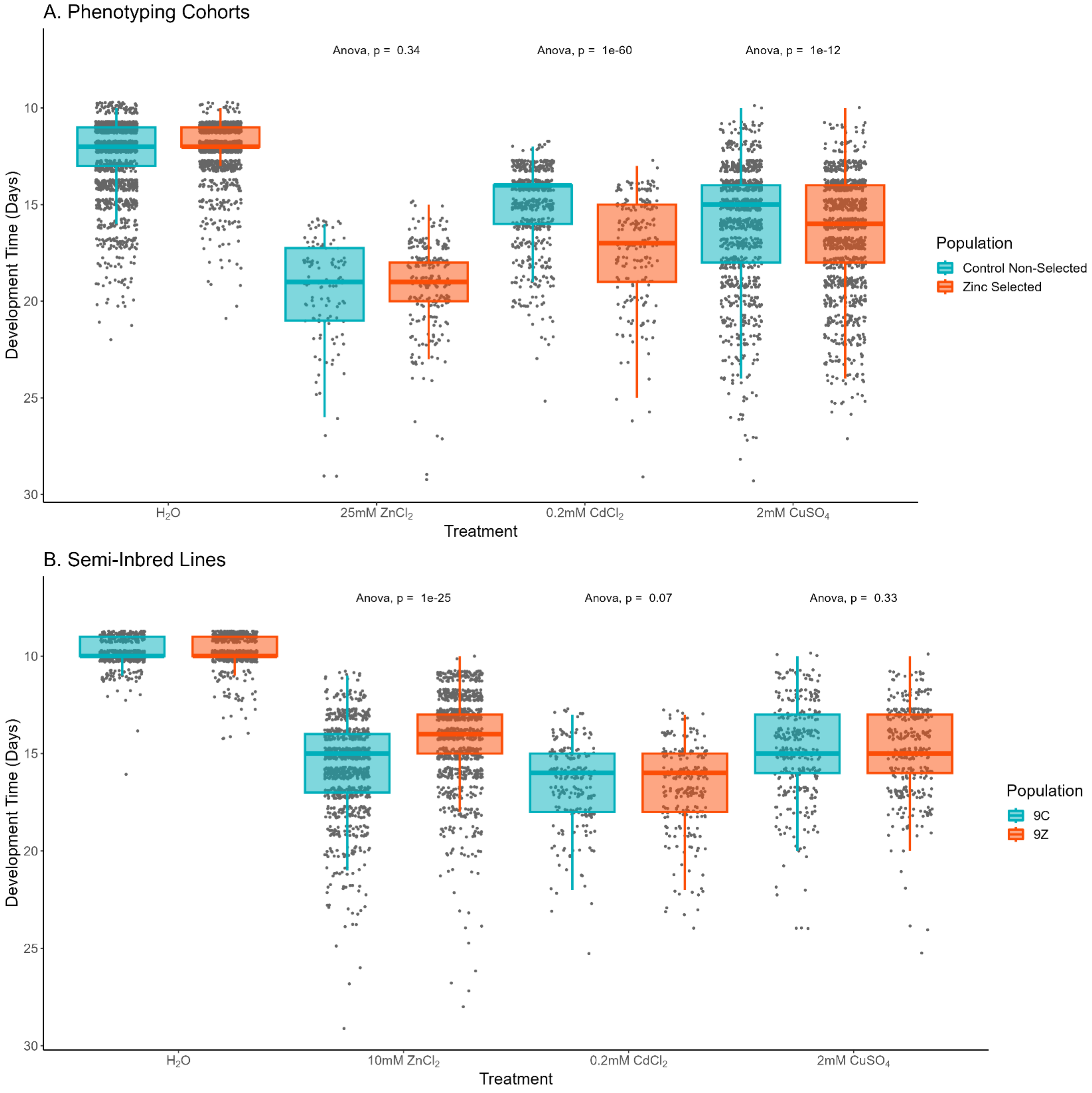
